# Estrogenic action by tris(2,6-dimethylphenyl) phosphate, an impurity in resorcinol bis[di(2,6-dimethylphenyl) phosphate] flame retardant formulations, impairs the development of female reproductive functions

**DOI:** 10.1101/539395

**Authors:** Kazuhiro Sano, Hidenori Matsukami, Go Suzuki, Nang Thinn Thinn Htike, Masahiro Morishita, Tin Tin Win Shwe, Shunji Hashimoto, Takaharu Kawashima, Tomohiko Isobe, Shoji F. Nakayama, Shinji Tsukahara, Fumihiko Maekawa

## Abstract

**Background:** Developmental exposure to environmental chemicals with estrogen-like activity has been suspected to permanently impair women’s health.

**Objectives:** In this study, we used a mouse model to evaluate whether a chemical having putative estrogen-like action detected by *in vitro* study, namely tris(2,6-dimethylphenyl) phosphate (TDMPP) impairs sexual differentiation of the brain.

**Methods:** To induce developmental exposure, TDMPP was administered subcutaneously to dams from gestational day 14 to parturition and to pups from postnatal day 0 to 9 at two different doses (TDMPP-high and TDMPP-low groups, respectively). To compare the results between TDMPP and typical estrogen exposures, 17β-estradiol was administered at two different doses on the same treatment schedule (E_2_-low and E_2_-high groups, respectively). A vehicle control group was formed by administering an equivalent volume of sesame oil to dams and to pups.

**Results:** Although there was no specific impairment in female ovary morphology, precocious puberty, detected by vaginal opening, and irregular estrous cycles, detected by vaginal cytology after sexual maturation, were found in TDMPP-and E_2_-treated groups, but not in the vehicle control group. In addition, lower lordosis response during reproductive behavioral tests was found in TDMPP-or E_2_-treated groups. To further clarify whether TDMPP directly affects sexual differentiation of the brain, we evaluated the transfer of TDMPP into the brain and the formation of sexual dimorphic nuclei. We detected a certain amount of TDMPP and its metabolites in the mouse brain after treatment, and masculinization of sexual dimorphic nuclei in the hypothalamus of female mice, suggesting the direct impact of TDMPP in developing brain.

**Discussion:** Taken together, the experimental evidence demonstrates that TDMPP directly enters the fetal and neonatal brain, inducing changes of sex-related brain structures, and impairing female reproductive functions.

## Introduction

One of the “Sustainable Development Goals” accepted by the United Nations in 2015 is gender equality (United Nations Statistical Commission, 2016). In the field of environmental health, gender-specific health issues related to sustainable development have gained increasing attention, and the inclusion of sex and gender differences in environmental health research has been proposed (Langer et al., 2015). Among environmental health problems related to sex and gender, sex difference of the biological responses when exposed to chemicals should be addressed more intensively, because the current male-bias in the use of experimental animals (Prendergast et al., 2014), even when evaluating the toxicity of new products, might prevent the detection of harmful effects in females. In particular, the effect of endocrine-disrupting chemicals has been known to be considerably sex-biased, and the effect of the exposition to endocrine disruptors in a vulnerable period such as the perinatal period is especially serious because of the permanence of the effects (Diamanti-Kandarakis et al., 2009). Therefore, the systematic evaluation of the impairment of sexual differentiation by new products using both males and females is required to elucidate their harmful effects.

In mammals, sexual differentiation of physiology and behavior during development is controlled not only by sex chromosomes, but also by gonadal steroid hormones. It has been reported that testosterone secreted from the testes during the developmental period in rodents is critical for determining the orientation of various kinds of behaviors, including sexual and social behaviors (Phoenix et al., 1959). In empirical studies using rodents, neonatal injection of testosterone was reported to cause abnormality in the estrous cycle, detectable by vaginal cytology, and to decrease female reproductive receptivity to males after sexual maturation, detectable by a typical receptive behavior called lordosis (Maekawa et al., 2014). Since the impairments in female-specific physiological and behavioral parameters have also occurred by neonatal injection of estrogen instead of testosterone, estrogen converted from testosterone in the brain by aromatase is thought to mediate them (MacLusky and Naftolin, 1981). Not only endogenous estrogen, but also chemicals with estrogenic activity, known as xenoestrogens, have been known to impair the development of female physiology and behavior. Estrogen affects sexual differentiation of the brain by acting on estrogen receptor alpha (ER-α), because the typical sexually dimorphic nuclei of the preoptic area (SDN-POA) and the bed nucleus of the stria terminals (BnST) were reported to develop in an ER-α dependent manner (Patchev et al., 2004; Tsukahara et al., 2011). Among xenoestrogens, bisphenol-A, used in the manufacturing of polycarbonate products, has become notorious, because a considerable amount of empirical evidence shows that it exerts its endocrine-disrupting action at least in part through ER-α (Gould et al., 1998), even if other estrogen receptors could also be related to its harmful effects (Alonso-Magdalena et al., 2012). More recently, the chemicals used for flame retardation, such as polybrominated diphenyl ethers (BDEs), have been reported to impair brain development by affecting thyroid-related and estrogenic cellular pathways (Zhou et al., 2002; Meerts et al., 2001). Therefore, the use of a certain type of penta-BDE and octa-BDE technical formulations such as BDE-47 and BDE-99 has been strictly regulated in many countries (European Parliament, 2002) and decabromodiphenyl ether was recently listed as persistent organic pollutants under the Stockholm Convention. Other than brominated flame retardants, phosphate flame retardants are used worldwide (Van der Veen et al., 2012). However, the evaluation of possible endocrine-disrupting actions of phosphate flame retardants is less advanced compared to that of brominated flame retardants. Recently, one of the co-authors of this study conducted a study demonstrating that tris(2,6-dimethylphenyl) phosphate (TDMPP), also known as 2,6-TXP, exerts estrogenic action at a level corresponding to about 1/10,000 of that of estradiol, by *in vitro* reporter assay (Suzuki et al, 2013). Since TDMPP is an impurity in flame retardant formulations of resorcinol bis[di(2,6-dimethylphenyl) phosphate] (PBDMPP) (Matsukami et al. 2015), whose demand has been increasing by the repressive trend of usage of BDEs, the possibility to be environmentally exposed to TDMPP will predictably increase. On the other hand, no study has been conducted whether TDMPP reveals endocrine-disrupting action *in vivo*.

In this study, we evaluated whether TDMPP impairs sexual differentiation of the brain using a mouse model. Moreover, to clarify whether this compound directly enters the developing brain, we measured the level of TDMPP transferred to the brain after maternal and neonatal injection. From the results of our toxicological and exposure studies, we determined that TDMPP is a novel endocrine disruptor acting directly on the mammalian brain.

## Methods

### Animals and developmental exposure toTDMPP

Pregnant C57BL/6J dams purchased from CLEA Japan (Tokyo, Japan) were used for perinatal exposure to TDMPP. The day on which a vaginal plug was detected was defined as gestational day (GD) 0. We prepared two experimental groups with developmental exposure to TDMPP at different doses, to discover impairments arising when perinatal mice were treated with TDMPP throughout the critical period of brain sexual differentiation. From GD 14 to parturition, TDMPP (99.9%, Hayashi Pure Chemical Ind., Ltd., Osaka, Japan), dissolved in sesame oil at the dose of 500µg/0.2 ml sesame oil/day for the TDMPP-low dose group, and 5,000 µg/0.2 ml sesame oil/day for the TDMPP-high dose group, was subcutaneously administered to dams. On top of the prenatal exposure, pups from postnatal day (PND) 0 to 9 were subcutaneously administered TDMPP at a dose of 50 µg/20 µl sesame oil/day for the TDMPP-low group and 500 µg/20 µl sesame oil/day for the TDMPP-high group. A vehicle control group (Oil group) was formed by administering an equivalent volume of sesame oil to dams and pups on the same experimental schedule. To reduce stress during treatments, we measured maternal and fetal body weight only at GD16 and PND0, respectively. Thus, by using the body weights measured, we estimated daily exposure levels of TDMPP-low and high dose groups. The maternal and fetal exposure level of TDMPP-low group was estimated to be 15 mg/kg bw/day and 38 mg/kg bw/day, respectively, and those of TDMPP-high group was estimated to be 146 mg/kg bw/day and 384 mg/kg bw/day, respectively. To compare the effects of TDMPP exposure to those of estrogen exposure, positive control groups were established by administering 17β-estradiol (E_2_, ≥ 98%, Sigma-Aldrich, St. Louis, MO, USA) dissolved in sesame oil at the dose of 0.5 µg/0.2 ml sesame oil/day for the E_2_-low group and 2 µg/0.2 ml sesame oil/day for the E_2_-high group, by subcutaneous injections to dams from GD 14 to parturition. As for TDMPP exposure, subcutaneous injections to pups from PND 0 to 9 were performed at the dose of 0.05 µg/20 µl sesame oil/day for the E_2_-low group, and 0.2 µg/20 µl sesame oil/day for the E_2_-high group. The doses of E_2_ were determined based on a previous report examining the relative action of TDMPP compared to estrogen on ER-α by *in vitro* CALUX assay (Suzuki et al., 2013): The action of the dose used in the E_2_-low group should be theoretically equivalent to the action of the dose used for the TDMPP-high group. Thus, the relative level of putative estrogenic action on ER-α in the five groups is the following: E_2_-high > E_2_-low ∼ TDMPP-high > TDMPP-low > Oil (vehicle control). Litters were weaned from their mothers on PND 21 and housed with same-sex littermates. Throughout the study, mice were housed in a room maintained at constant temperature (24 ± 1°C) and humidity (50 ± 10%) with a 12/12-h light/dark cycle. Food and water were provided *ad libitum*. The administration of TDMPP to sexually mature females was also performed in order to investigate whether TDMPP affects sexual receptivity in adults. The relevant methods and results are described in the Supplemental methods and results.

All procedures were approved by the Animal Care and Use Committee at the NIES and conducted in strict accordance with the NIES guidelines. All efforts were made to minimize the number of animals and their suffering.

### Transfer to brain

To determine the transfer of TDMPP to the fetal and neonatal brain, pregnant C57BL/6J females were purchased from CLEA Japan (Tokyo, Japan). Fifteen dams were subcutaneously injected with TDMPP (5,000 µg/0.2 ml sesame oil) on GD 16. Dams were sacrificed by decapitation and the brain and blood of the dams and the fetuses (1 to 3 of each sex per dam) were collected at the time points of 0, 8, 16, 24, and 48 h after injection (3 dams per time point). Pups born to five dams were subcutaneously injected with TDMPP (500 µg/20 µl sesame oil) on PND 1 and sacrificed by decapitation at the time points of 0, 8, 16, 24, and 48 h after injection (3 males from 3 dams and 3 females from 3 dams per time point), and brains were collected. Samples were immediately frozen in dry-ice and stored at −75 °C.

### Examination of general reproductive physiology and histology

After birth, all the pups were subjected to body weight (BW) and anogenital distance (AGD) measurements. The body weight was also measured at the time of weaning (PND 21) and at 10 weeks of age. The AGD was also measured at the time of weaning. All females were inspected daily for their first vaginal opening starting from PND 18 until the opening was observed. Vaginal smears were taken daily for 26 days starting from 9 weeks of age. Vaginal lavages were collected using 10 µl pipette tips thinly wrapped with cotton moistened with deionized water. The lavages were placed on a slide glass, air-dried and stained with 0.1% methylene blue solution. The estrous stage of each individual on each day was determined based on the criteria described in Cora et al. (2015), in which the estrous cycle is divided into 5 stages as follows: proestrus (P), estrus (E), metestrus-1 (M1), metestrus-2 (M2), and diestrus (D). When female mice reached 14 weeks of age, the ovaries were bilaterally removed under isoflurane anesthesia from selected mice and weighed. The ratio of ovarian weight to body weight was compared between groups. Ovary histology was examined by paraffin sectioning and conventional hematoxylin-eosin staining.

### Overall scheme of the behavioral test battery

When males and females reached 10 weeks and 14 weeks of age, respectively, one or two mice of each sex were randomly selected from each litter, separated from their littermates and housed individually in plastic cages (5 × 22 × 12 cm). At the time of isolation, selected females were ovariectomized under isoflurane anesthesia. They were subjected to a behavioral test battery for emotional and socio-sexual behaviors, consisting of open field test, light-dark transition test, and sexual behavior test for both sexes, and aggressive behavior test for males only. All behavioral tests were performed during the dark phase, starting more than 2 h after lights off. After completing the behavioral tests, mice were sacrificed and blood and brain samples were collected for enzyme immunoassays and immunohistochemistry. To eliminate the litter effect, data collected from individuals were first averaged within littermates of the same sex. Thus, the data shown in this study represent the mean value of the score per litter, unless otherwise specified. The experimental design is shown in Figure 1.

**Figure 1:**
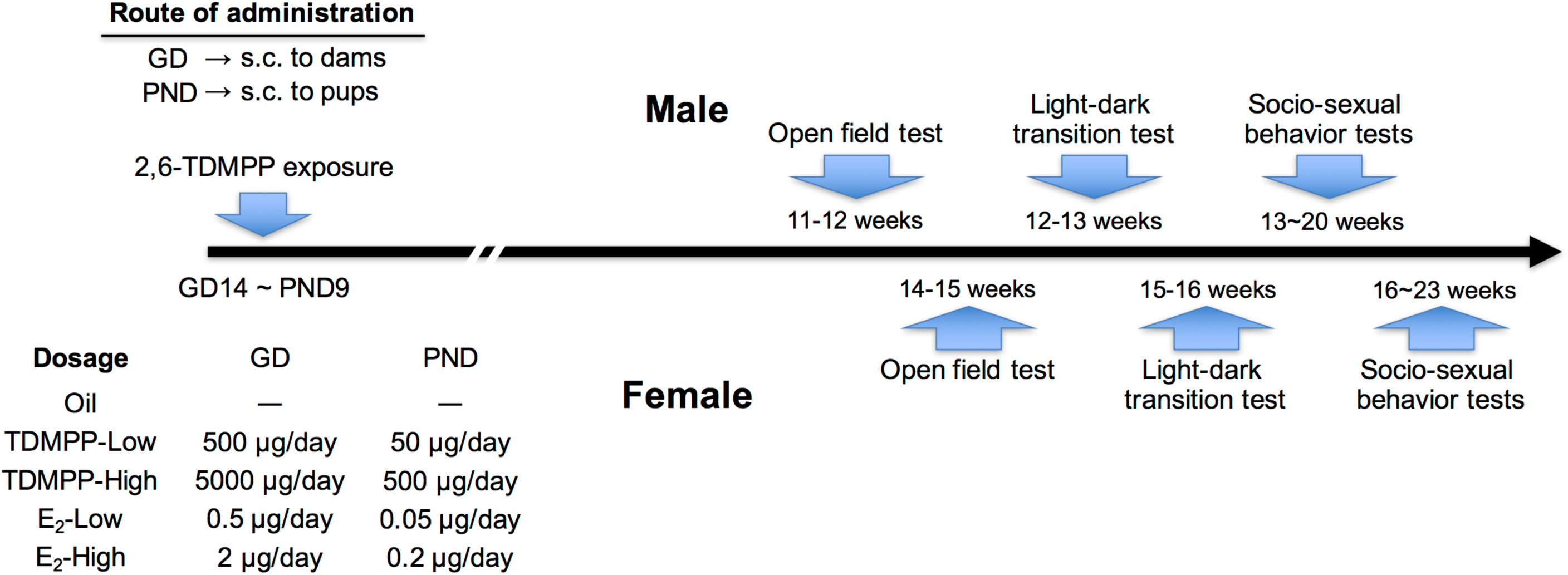
The experimental design of the developmental exposure and behavioral test battery.

### Open field test

Seven to 9 days after isolation, each mouse was tested in an open field apparatus (60 × 60 cm with 30 cm tall opaque walls) illuminated by white light (50 lux) twice in two consecutive days, each trial lasting 10 min. The floor of the apparatus was virtually divided into 25 square sections (12 × 12 cm each) and 9 inner squares were designated as the center area. At the beginning of each trial, the mouse was placed in a fixed corner, with the head facing the corner. Total moving distance (total distance) and time spent in the center area (center time) were measured digitally by an automated video tracking system (ANY-maze, Stoelting, USA).

### Light-dark transition test

Five to 7 days after the open field test, each mouse was tested in a light-dark transition test apparatus for 10 min. The test apparatus consisted of an enclosed dark and an open-top light compartment (30 × 30 × 30 cm each), connected by an inner doorway (3 × 3 cm) located in the center of the partition at the floor level. The open-top light compartment was brightly illuminated with white light (350 lux). At the beginning of the trial, the mouse was introduced in the dark compartment. The latency to enter the light compartment and the cumulative time spent in the light compartment were measured by an automated video tracking system (ANY-maze, Stoelting, USA) (Sano et al., 2016).

### Male socio-sexual behavior tests

Starting 5 to 7 days after the light-dark transition test, male mice were tested biweekly for sexual and aggressive behaviors. All tests were performed under red light illumination and videotaped. In the male sexual behavior test, each male was tested in its home cage for sexual behavior meant to lure female C57BL/6J mice. All lured females had been ovariectomized and primed with subcutaneous injections of estradiol benzoate (10 µg/0.1 ml dissolved in sesame oil) twice at 48 and 24 h before testing and progesterone (500 µg/ 0.1 ml dissolved in sesame oil) once at 4-6 h before testing to ensure high sexual receptivity. Three trials were performed, each lasting 30 min. The number of attempted mounts, mounts, and intromissions was scored for each mouse (Sano et al., 2016).

In the male aggressive behavior test, each mouse was tested in a resident-intruder paradigm against olfactory bulbectomized male C57BL/6J mice. On each trial week, the mice were tested on three consecutive days, for a total of 9 trials, each lasting 15 min. The number and duration of aggressive bouts toward the intruder were scored for each mouse. The data from the 3 trials performed each week were averaged for each mouse and used for statistical analysis. An aggressive bout was defined as a set of behavioral interactions that included at least one of the following actions: Chasing, boxing, wrestling, biting, tail-rattling, and offensive lateral attack. If the interval between 2 aggressive bouts did not exceed 3 seconds, the 2 bouts were considered to be continuous and scored as 1 bout (Sano et al., 2016).

### Female sexual behavior test

Five to 7 days after the light-dark transition test, female mice were tested for sexual behavior toward a sexually experienced ICR/JCL male mouse in the male’s home cage. The test was performed weekly for a total of 5 trials. Female mice were tested under an artificial estrous condition in which they had been primed with subcutaneous injection of estradiol benzoate (5 µg dissolved in 0.1 ml sesame oil) twice at 48 and 24 h before testing, and progesterone (250 µg dissolved in 0.1 ml sesame oil) once at 4-6 h before testing. Each test lasted until females received either 15 mounts or 15 intromissions. The number of lordosis responses to either mount or intromission was scored for each mouse. A lordosis quotient was calculated by dividing the number of lordosis responses by the 15 mounts or intromissions (Sano et al., 2016).

### Blood and brain sampling

After the completion of the behavioral testing, mice were deeply anesthetized with a solution of a 1:1 mixture of sodium pentobarbital (60 mg/kg BW) and heparin (1,000 units/ml), and blood was collected from the left ventricle. They were then transcardially perfused with 0.1 M phosphate-buffer (PBS; pH 7.2), followed by 4% paraformaldehyde (PFA) in 0.1 M PBS. Brains were removed, post-fixed with 4% PFA in 0.1 M PBS overnight at 4°C, and cryoprotected in 0.1 M PBS containing 30% sucrose. Sections (30 µm thick) were made on a freezing microtome (REM-710, Yamato, Japan) at 120 µm intervals. Male and female brains of the Oil, TDMPP-low, TDMPP-high and E_2_-low groups were used for immunohistochemistry to detect sexually dimorphic nuclei.

### Enzyme immunoassay for testosterone and estradiol

Samples were extracted from male plasma (100 µl) with ethyl acetate. Testosterone and estradiol concentrations were determined using enzyme immunoassay kits for each hormone (Cayman Chemicals, Ann Arbor, MI, USA) according to the manufacturer’s instructions.

### Calbindin D-28K immunohistochemistry to detect sexual dimorphic nuclei

Endogenous peroxidase in the brain section was removed by incubating with 0.6% H_2_O_2_ containing 0.05 M PBS with 1% Triton X-100 (PBST) for 60 min at room temperature. The sections were treated with 5% normal goat serum containing PBST for 60 min at room temperature to prevent the nonspecific binding of the antibody. Afterward, they were incubated with a monoclonal antibody against CB (C9848, Sigma Aldrich, St. Louis, MO, USA, 1:15,000) for 2 days at 4°C. Subsequently, the sections were rinsed with PBST and incubated in a peroxidase-labeled polymer conjugated with goat anti-mouse immunoglobulin (Dako Envision Plus, Dako, Carpinteria, CA, USA) for 30 min at room temperature. After rinsing again with PBST, the sections were stained with 3,3′-diaminobenzidine in chromogen solution (Dako). Finally, they were mounted on gelatin-coated slides.

### Delineation of sexually dimorphic nuclei

CB is a protein maker for detecting two specific sexually dimorphic nuclei: The calbindin-sexually dimorphic nucleus (Calb-SDN), subregion of the medial preoptic area; and the principal nucleus of the bed nuclei of the stria terminalis (BNSTp) (Budefeld et al. 2008, Gilmore et al. 2012, Orikasa & Sakuma 2010, Sickel & McCarthy 2000). We defined the Calb-SDN as the distinctive ellipsoidal cluster of CB-immunoreactive cells at the preoptic area/anterior hypothalamus, dorsolaterally angled from the third ventricle, and located dorsal to the optic chiasm, lateral to the third ventricle, and ventral to the BNSTp (Gilmore et al, 2012). We also defined the BNSTp as the clusters of CB-immunoreactive cells between the stria terminalis and the stria medullaris of the thalamus, in the area surrounded by the lateral ventricle and the third ventricle (Gilmore et al. 2012, Moe et al. 2016, Wittmann & Mclennan 2013).

### Stereological analysis of sexually dimorphic nuclei

The CB-stained sections were observed under the light microscope. The volume and number of CB-immunoreactive (CB-ir) cells in Calb-SDN and BNSTp were analyzed using the Stereo Investigator software (MBF Bioscience Inc., Williston, VT, USA). Since each slide was randomly assigned an identification number not related to the original number of the animal, the observer who performed the analysis was blinded to the sample origin. The optical fractionator method of the stereological probe workflow in the software was used to analyze the CB-stained sections. The outlines of Calb-SDN and BNSTp were traced on the left side of brain sections to determine the analysis area according to a mouse brain atlas (Paxinos & Franklin, 2004). The CB-ir cells were counted in a defined counting frame and grid for each area. Details on the analysis are reported in Table 1.

**Table 1:** Parameters for stereological analyses for Calb-SDN

| 4 |  | <u>Oil</u> |  | <u>TDMPP-low</u> |  | <u>TDMPP-high</u> |  | <u>E2</u> |  |
| --- | --- | --- | --- | --- | --- | --- | --- | --- | --- |
| 5 |  | Male | Female | Male | Female | Male | Female | Male | Female |
| 6 | No. of sections | 9.8±0.67 | 4.2±0.2 | 10.3±0.33 | 7.6±0.51 | 9±0.32 | 7.6±0.68 | 9.2±0.2 | 7.2±0.37 |
| 7 | Sampling Grid Size | 70×70 µm |  |  |  |  |  |  |  |
| 8 | Counting frame size | 35×35 µm |  |  |  |  |  |  |  |
| 9 | Number of sampling sites | 51.8 ± 4.22 | 20.4 ± 1.21 | 98.7 ± 4.91 | 48 ± 4.16 | 68.2 ± 3.85 | 46.6 ± 4.77 | 64.6 ± 4.76 | 41.4 ± 2.98 |

**Parameters for stereological analyses for BNSTp**
| 13 |  | <u>Oil</u> |  | <u>TDMPP-low</u> |  | <u>TDMPP-high</u> |  | <u>E2</u> |  |
| --- | --- | --- | --- | --- | --- | --- | --- | --- | --- |
| 14 |  | Male | Female | Male | Female | Male | Female | Male | Female |
| 15 | No. of sections | 10.8±0.37 | 9±0.45 | 10.7 ± 0.33 | 9.4 ± 0.4 | 11.2 ± 0.37 | 9.6 ± 0.24 | 10.8±0.2 | 9.8±0.37 |
| 16 | Sampling Grid Size | 100 × 100 µm |  |  |  |  |  |  |  |
| 17 | Counting frame size | 50 × 50 µm |  |  |  |  |  |  |  |
| 18 | Number of sampling sites | 227 ± 9.9 | 168 ± 11.53 | 257.7 ± 9.17 | 202.7 ± 3.53 | 226.8 ± 9.30 | 190 ± 3.65 | 245 ± 14.24 | 202.6 ± 7.28 |

19 Section thickness, 30 µm, section interval, 30 µm; dissector height 14-16 µm; guard zone height, 1.5 µm

### Chemical analysis of TDMPP and its metabolites in brain

After thawing of the brain samples, tris(3,5-dimethylphenyl-d9) phosphate (Hayashi Pure Chemical Ind., Ltd.) was added as an internal standard, and the samples were homogenized by an ultrasonic homogenization device and extracted with methanol. The crude extract was passed through the Oasis Wax (150 mg/30 µm) cartridge column (prewashed with 5 ml of methanol). Two fractions were collected: 3 mL of methanol (fraction 1) and 5 mL of 0.5% ammonium hydroxide in methanol (fraction 2). The eluate of fraction 1 was passed through the ENVI-Carb 250 mg cartridge column (prewashed with 5 ml of dichloromethane:toluene (3:1, v/v) mixture). 3 ml of dichloromethane:toluene (3:1, v/v) mixture was passed through the ENVI-Carb column and the eluate was collected (fraction 1A). The eluates of fraction 1A and 2 were evaporated and redissolved in 0.5 ml of methanol. An electrospray ionization-quadrupole time-of-flight mass spectrometer (ESI-QTOF-MS) equipped with an ultra-high-performance liquid chromatograph (LC) system (1290 Infinity/6530 Accurate-Mass QTOF LC/MS system; Agilent Technologies Inc., Santa Clara, CA, USA) with a reversed-phase LC column (ZORBAX Eclipse Plus C18 RRHD, 50 mm × 2.1 mm i.d., 1.8 mm; Agilent Technologies Inc.) was used for the identification and quantification of TDMPP and its metabolites. The mass range for the MS investigation was set at m/z 100−1500. The mass range for MS/MS investigation was set to m/z 50−1000. The inter-and intra-day variation of measurements were 14% and 12%, respectively, and the mean recovery rate of internal standard from fetal and neonatal brains was 75%.

### Statistical Analysis

All data are presented as the mean ± standard error of the mean (SEM). Data were analyzed using analysis of variance (ANOVA) followed by Bonferroni *post hoc* tests or Student t-test. Differences were considered statistically significant at p-values less than 0.05. The analysis was performed with the SPSS 19.0 statistical package (SPSS Inc., Chicago, IL, USA).

## Results

### Body weight

No statistically significant effect of TDMPP exposure on body weight of either males or females at birth, PND 21, or 10 weeks of age was found (Figure 2A-F). In addition, there was no significant difference in body weight between the control Oil group and the E_2_-treated groups, whereas a slight but significant difference was found among the E_2_-treated groups. A detailed description of the statistical analysis is reported in the Supplementary Results.

**Figure 2:**
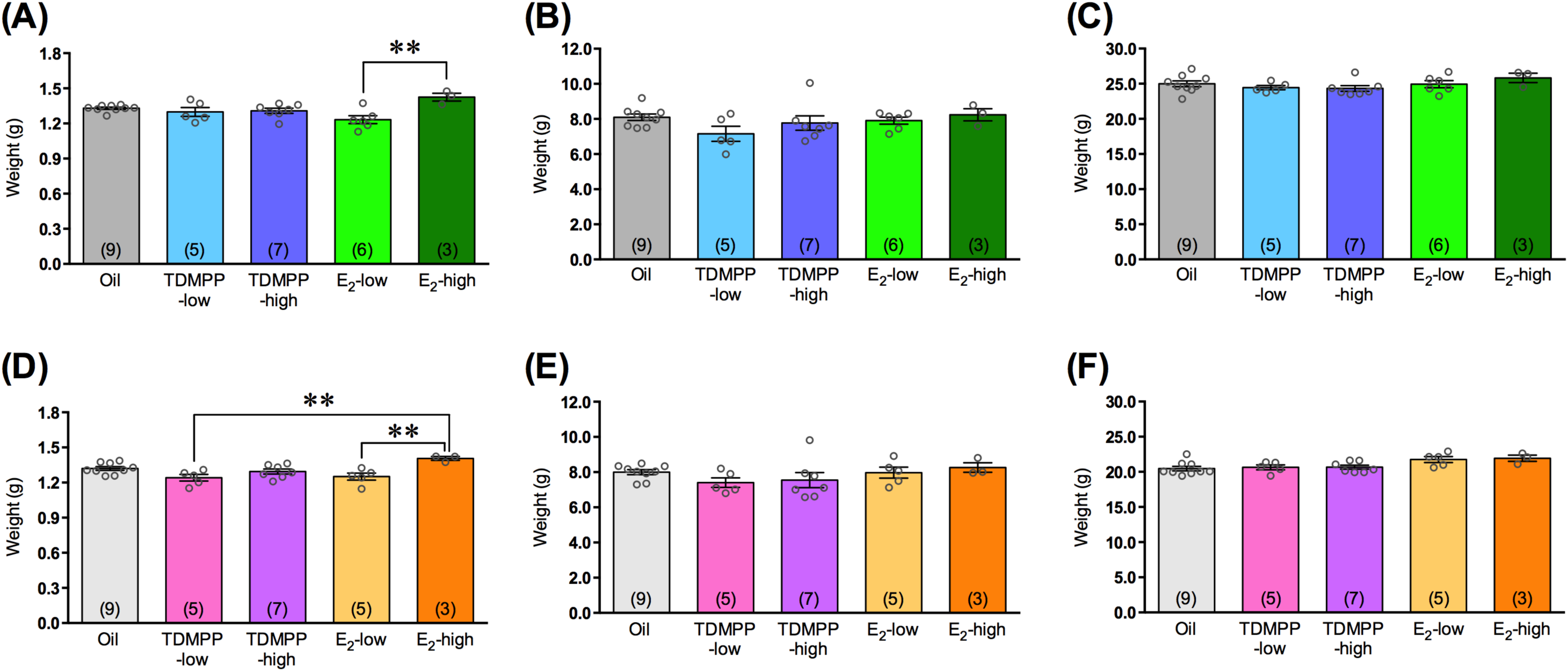
Effect of developmental TDMPP exposure on body weight. **(A-C)** male and **(D-F)** female mice of each treatment group. The body weight at birth **(A,D)**, PND 21 **(B,E)**, and 10 weeks of age **(C,F)**. The number in parentheses indicate number of liters. The data are presented as the mean ± SEM. \*\**P* < 0.01.

### Anogenital distance

A statistically significant effect of TDMPP exposure on AGD was found by ANOVA among the male groups at birth and PND 21, but we could not detect which groups caused the significant difference, at least by Bonferroni *post hoc* test (Figure 3A-D). There was no significant difference among the female groups at birth and PND 21. A detailed description of the statistical analysis is reported in the Supplemental results.

**Figure 3:**
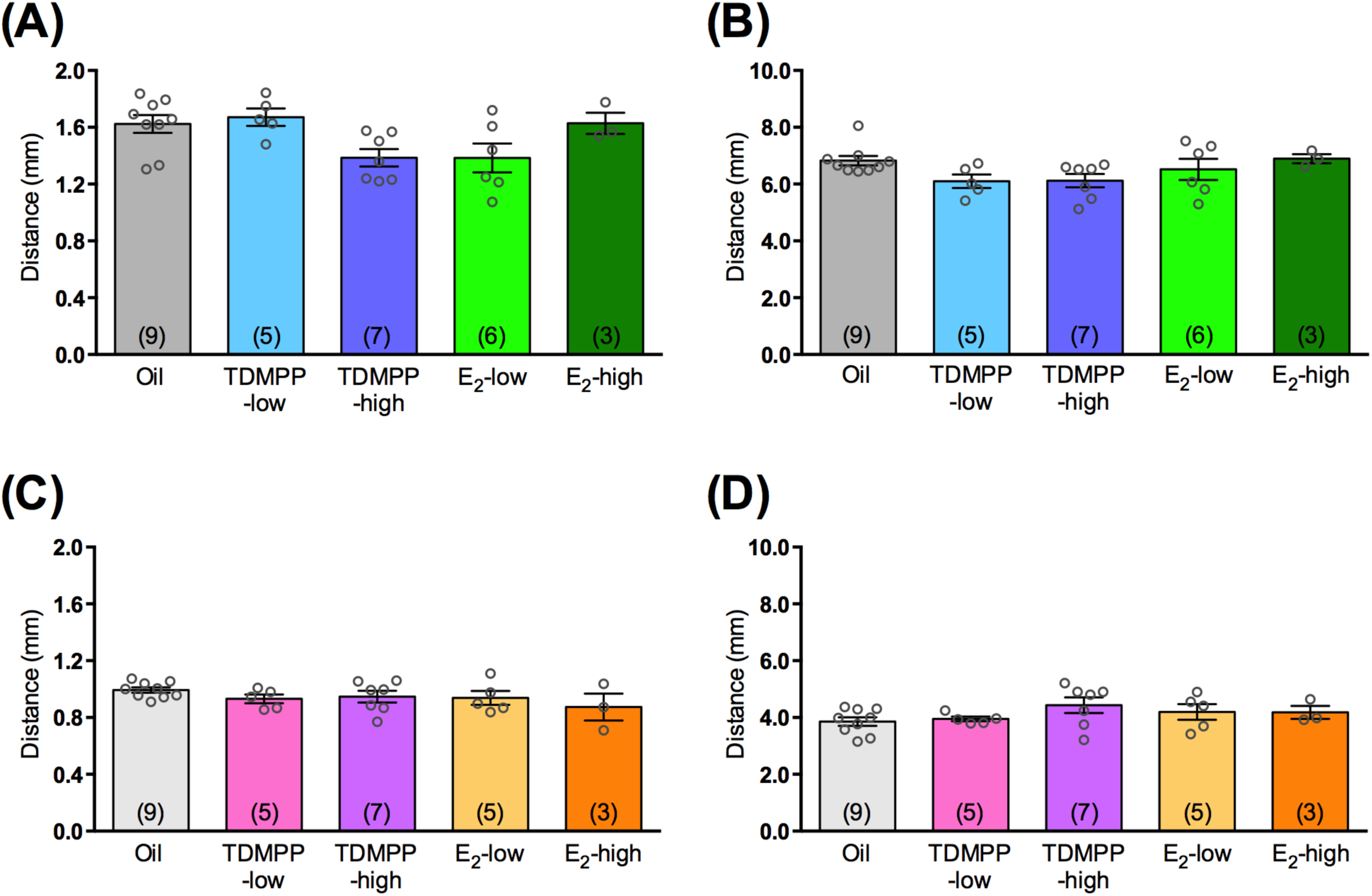
Effect of developmental TDMPP exposure on anogenital distance. **(A,B)** male and **(C,D)** female of each treatment group. The anogenital distance at birth **(A,C)** and PND 21 **(B,D)**. The number in parentheses indicate number of liters. The data are presented as the mean ± SEM.

### Female reproductive physiology and histology

In the TDMPP-high and E_2_-high groups, the day we first observed vaginal opening was significantly earlier compared to the Oil, E_2_-low, and TDMPP-low groups [*F*(4,17) = 20.132, *P* < 0.001; Bonferroni *post hoc* test: *P* < 0.001, TDMPP-high and E_2_-high vs. Oil or TDMPP-low; *P* = 0.011, TDMPP-high vs. E_2_-low; *P* = 0.018, E_2_-high vs. E_2_-low; Figure 4A,B]. The estrous cycle, detected by vaginal smear after sexual maturation, was impaired in both the TDMPP and E_2_-exposed groups (Figure 5A-E). The number of estrous days in the TDMPP-low, TDMPP-high, E_2_-low and E_2_-high groups was significantly reduced compared to the Oil group [*F*(4,24) = 25.228, *P* < 0.001; Bonferroni *post hoc* test: *P* < 0.001, TDMPP-high, E_2_-low, and E_2_-high vs. Oil; *P* = 0.011, TDMPP-low vs. Oil; Figure 5F]. No difference was found in the ratio of ovarian weight to body weight between the groups [*F*(4,24) = 2.211, *P*= 0.098; Figure 6].

**Figure 4:**
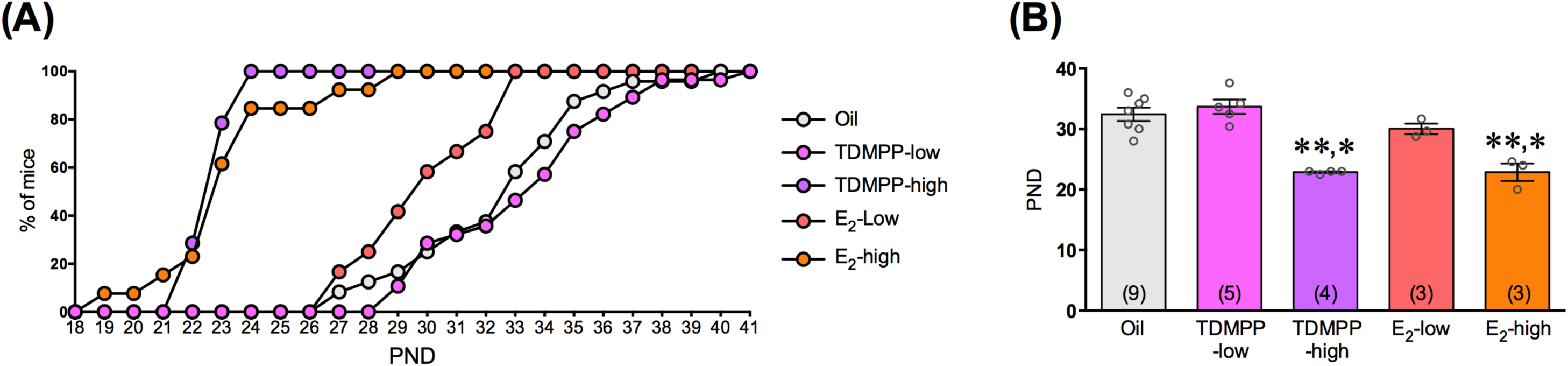
Effect of developmental TDMPP exposure on first vaginal opening. **(A)** Percentage of females displayed vaginal opening. (B) Average ages of first vaginal opening. The number in parentheses indicate number of liters. The data are presented as the mean ± SEM. \*\**P* < 0.01 vs Oil or TDMPP-low. \**P* < 0.05 vs E_2_-low.

**Figure 5:**
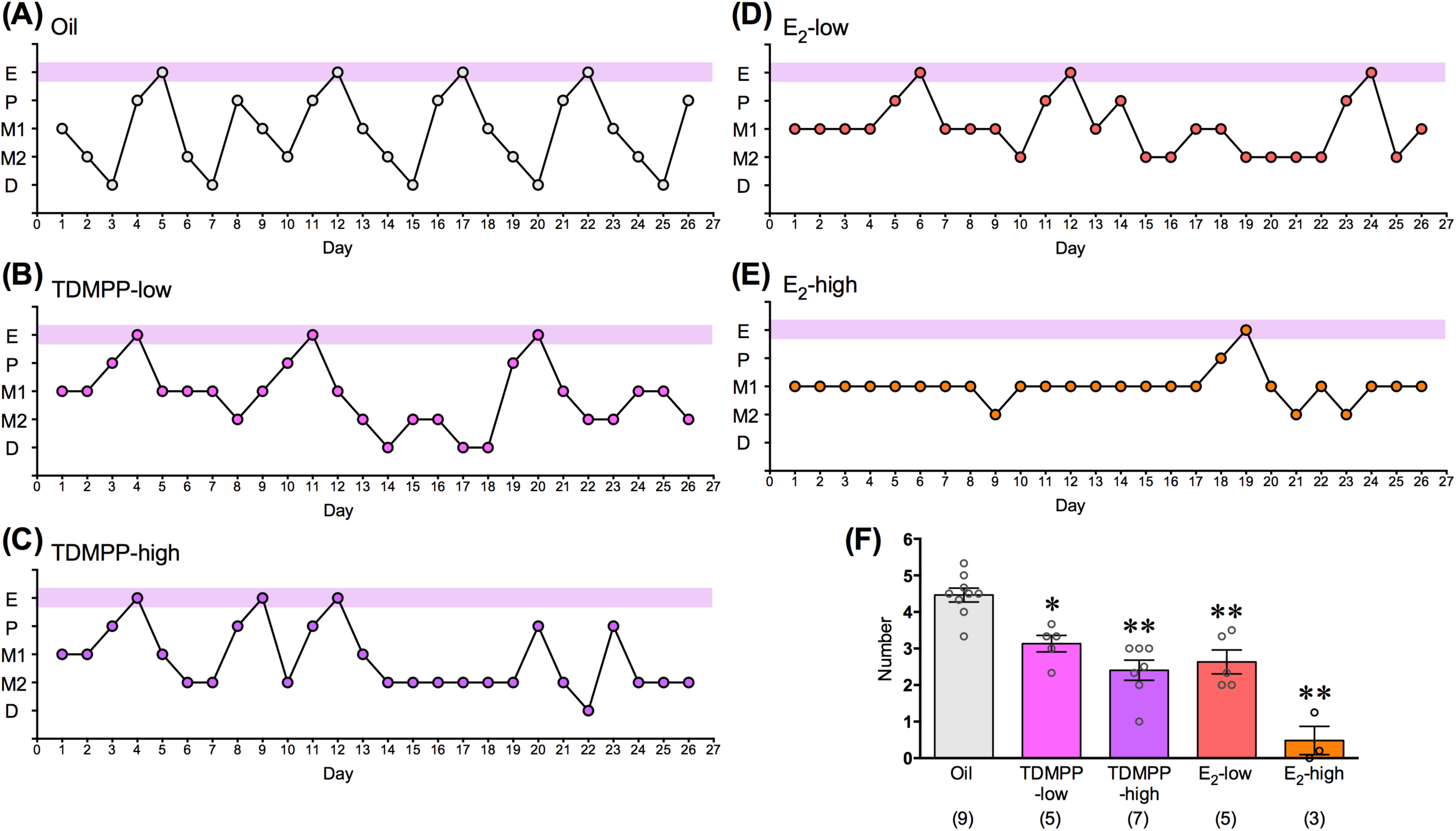
Effect of developmental TDMPP exposure on estrous cycle. **(A-E)** The representative estrous cycle pattern of each treatment group. E = estrus, P = proestrus, M1 = metestrus 1, M2 = metestrus 2, D = diestrus. **(F)** The number of cycles within the 27 days of recording period. The number in parentheses indicate number of liters. The data are presented as the mean ± SEM. \*\**P* < 0.01, \**P* < 0.05 vs Oil.

**Figure 6:**
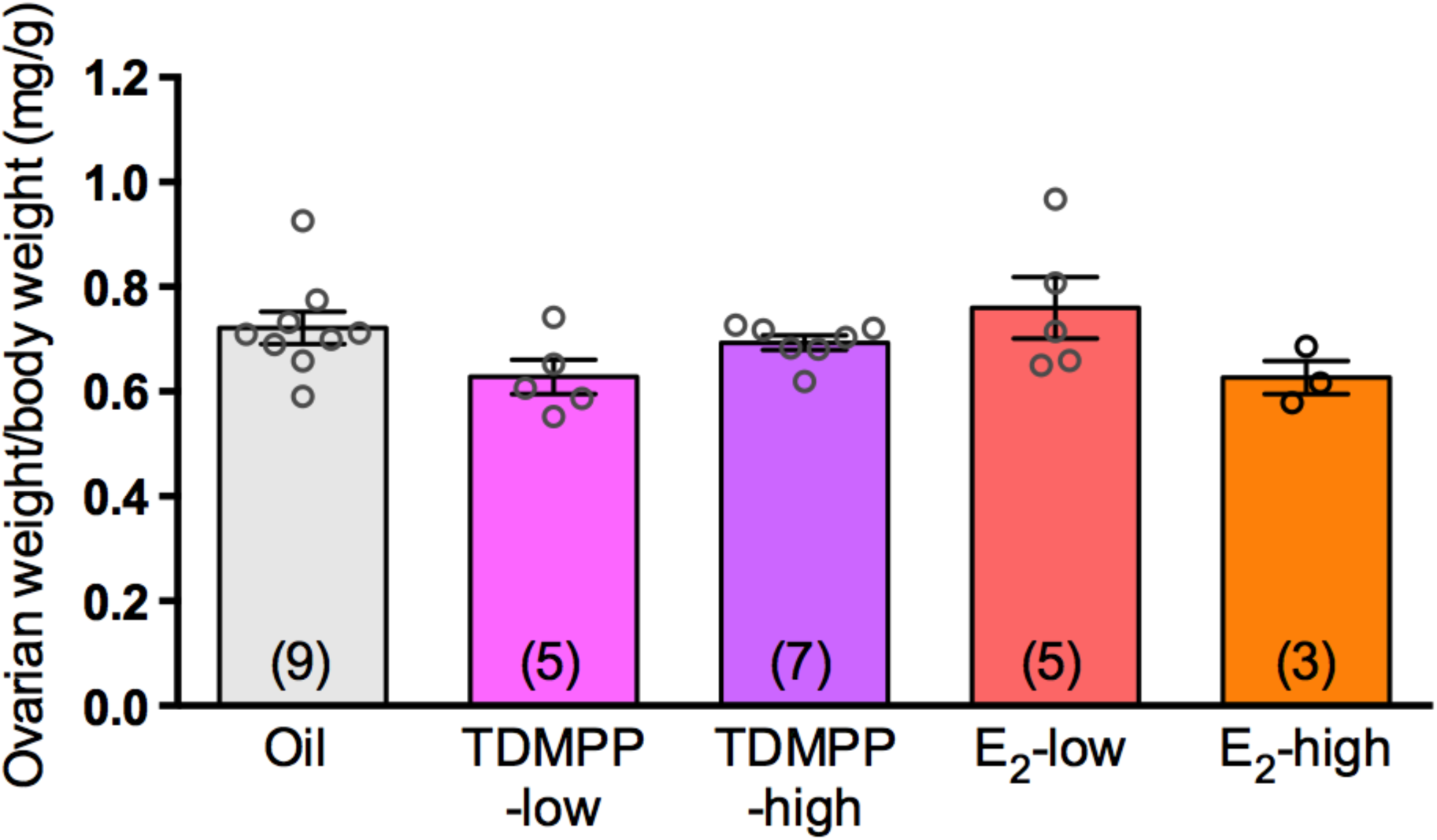
Effect of developmental TDMPP exposure on the ratio of ovarian weight to body weight. The number in parentheses indicate number of liters. The data are presented as the mean ± SEM.

### Open field test

In males, no difference was found in total moving distance or center time between the groups [Figure 7A and B]. In females, there was no significant difference in total moving distance or center time between the Oil group and other groups, whereas there was a significant difference in total moving distance between the E_2_-treated groups and the TDMPP-low group [Figure 7C and D]. A detailed description of the statistical analysis is reported in the Supplemental results.

**Figure 7:**
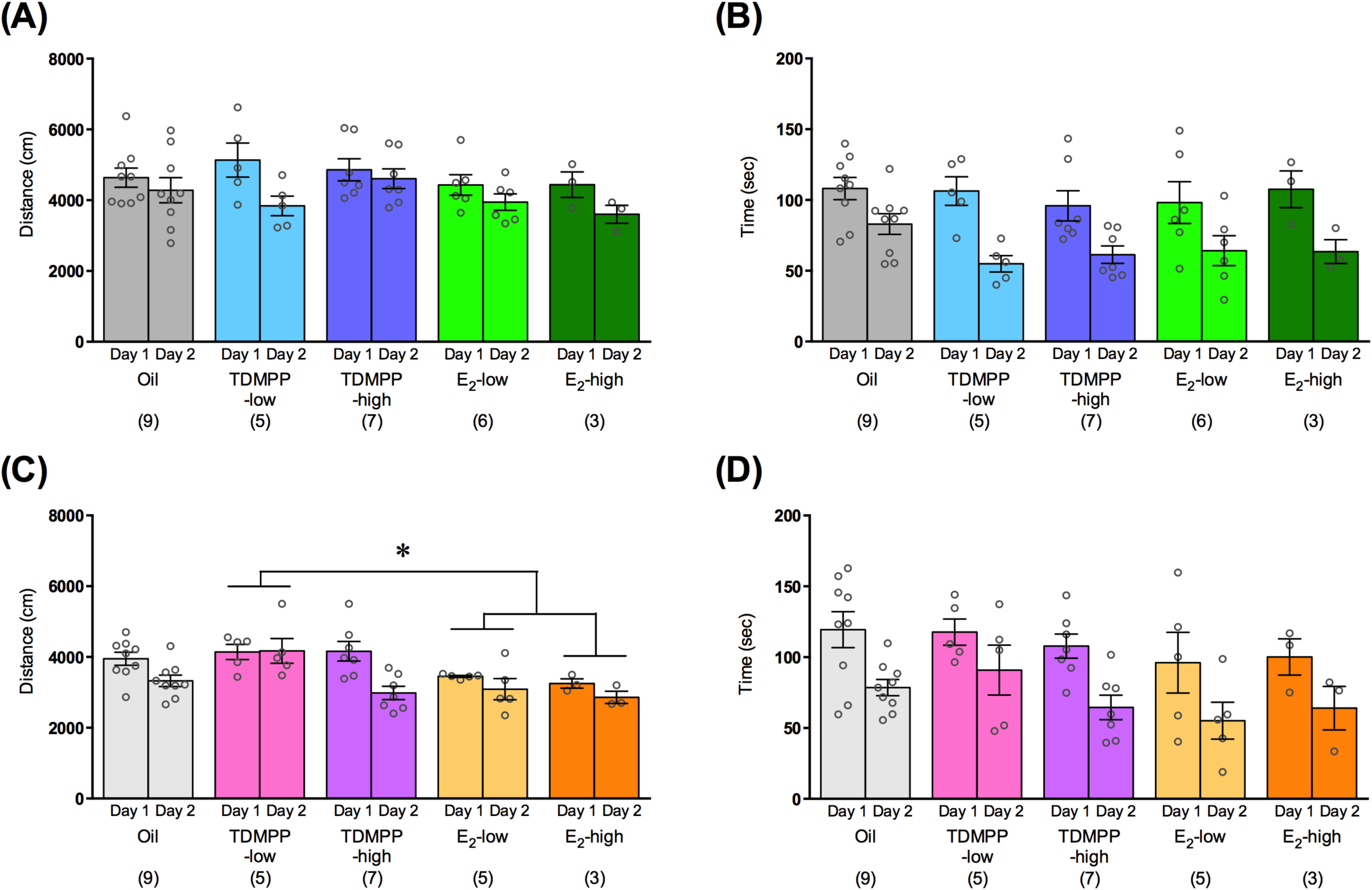
Effect of developmental TDMPP exposure on the open field activity. **(A,B)** male and **(C,D)** female of each treatment group. **(A,C)** the total moving distance and **(B,D)** time spent in center area of open field test apparatus. The number in parentheses indicate number of liters. The data are presented as the mean ± SEM.

### Light-dark transition test

There was no difference in the time spent in the light compartment between the Oil group and the TDMPP-treated groups in males, whereas the male mice in the E_2_-high group spent significantly shorter time in the light compartment than the mice in other male groups [*F*(4,25) = 7.672, *P* < 0.001; Bonferroni *post hoc* test: *P* = 0.001, E_2_-high vs. Oil or TDMPP-high; *P* = 0.002, E_2_-high vs. E_2_-low; *P* = 0.030, E_2_-high vs. TDMPP-low; Figure 8A]. The latency to enter the light compartment did not differ among the groups [*F*(4,25) = 1.739, P = 0.173; Figure 8B]. In females, no differences were found among the groups in the time spent in the light compartment [*F*(4,24) = 2.292, *P* = 0.089; Figure 8C] or the latency to enter the light compartment [*F*(4,24) = 0.306, *P* = 0.871; Figure 8D].

**Figure 8:**
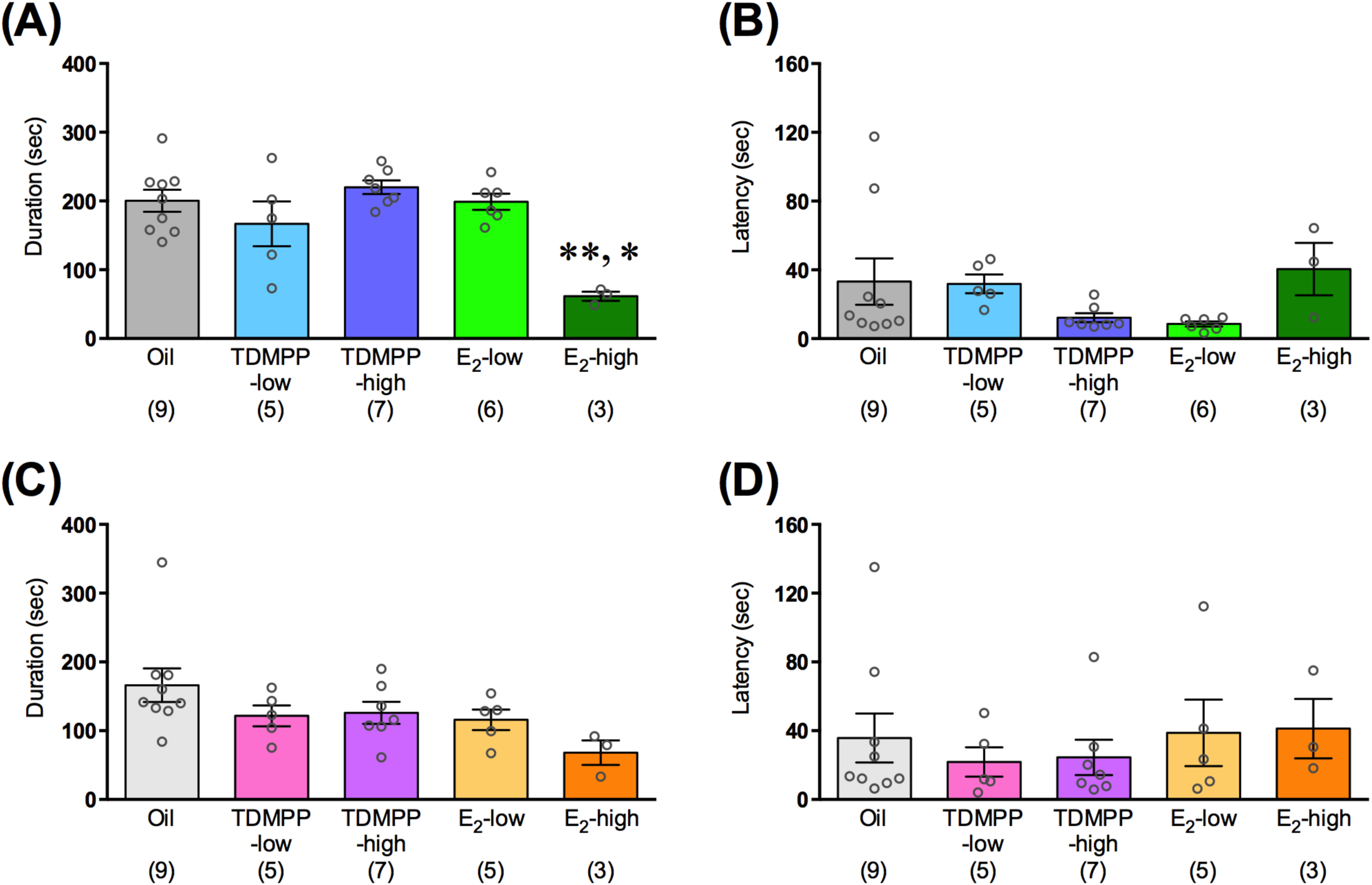
Effect of developmental TDMPP exposure on anxiety-related behavior, as measured in the light-dark transition test. **(A,B)** male and **(C,D)** female of each treatment group. (A,C) the total time spent in, and (B,E) the latency to enter the light compartment of the light-dark transition apparatus. The number in parentheses indicate number of liters. The data are presented as the mean ± SEM. \*\**P* < 0.01 vs Oil, TDMPP-high, or E_2_-low, \**P* < 0.05 vs TDMPP-low.

### Male socio-sexual behavior tests

Regarding male sexual behavior, no differences between the Oil group and the TDMPP-treated groups were found in the number of attempted mounts [treatment, *F*(4,25) = 0.476, *P* = 0.753; treatment × test number, *F*(8,50) = 1.259, *P* = 0.286; Figure 9A], mounts and intromissions [treatment, *F*(4,25) = 1.226, *P* = 0.325; treatment × test number, *F*(8,50) = 0.441, *P* = 0.890; Figure 9C]. No differences were found between the Oil group and the E_2_-treated groups either, although the mice in the E_2_-high group showed significantly higher number of mounts compared to the E_2_-low group [treatment: *F*(4,25) = 2.831, *P* = 0.046; Bonferroni *post hoc* test: *P* = 0.050, E_2_-high vs. E_2_-low; treatment × day: *F*(4,25) = 1.519, *P* = 0.174; Figure 9B]. As for aggressive behavior, the mice in the TDMPP-high group showed significantly higher number of aggressive bouts compared to the Oil, TDMPP-low and E_2_-low groups [treatment: *F*(4,25) = 6.403, *P* = 0.001; Bonferroni *post hoc* test: *P* = 0.005, TDMPP-high vs. Oil; *P* = 0.002, TDMPP-high vs. TDMPP-low; *P* = 0.030, TDMPP-high vs. E_2_-low; treatment × day: *F*(8,50) = 0.575, *P* = 0.793; Figure 10A]. A similar tendency was observed in the total duration of aggressive bouts, but the difference was not statistically significant [treatment: *F*(4,25) = 2.158, *P* = 0.103; treatment × day: *F*(8,50) = 1.147, *P* = 0.350; Figure 10B].

**Figure 9:**
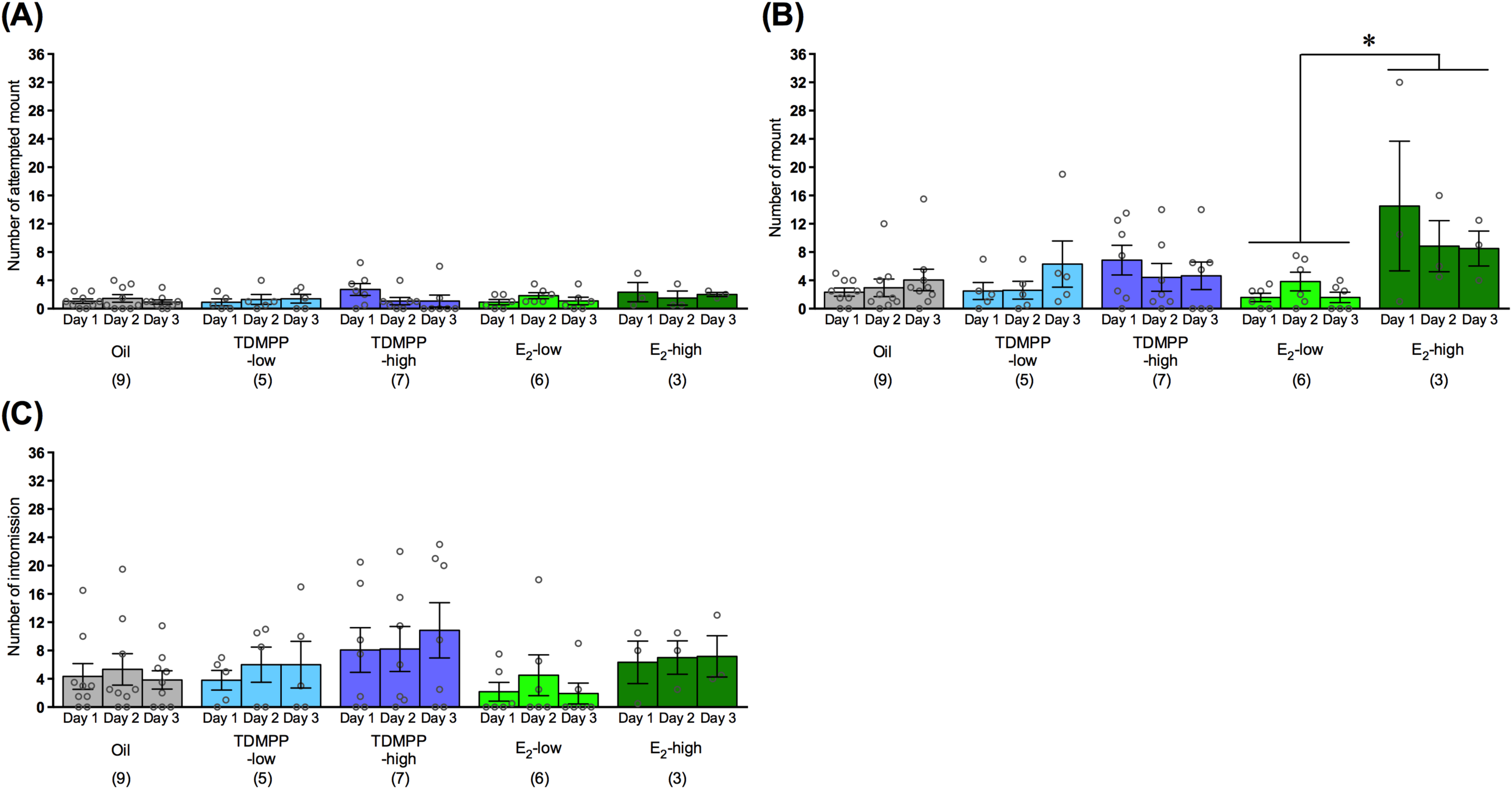
Effect of developmental TDMPP exposure on male sexual behavior. The total number of **(A)** attempted mount, **(B)** mount, and **(C)** intromission toward female stimuli. The number in parentheses indicate number of liters. The data are presented as the mean ± SEM. \**P* < 0.05.

**Figure 10:**
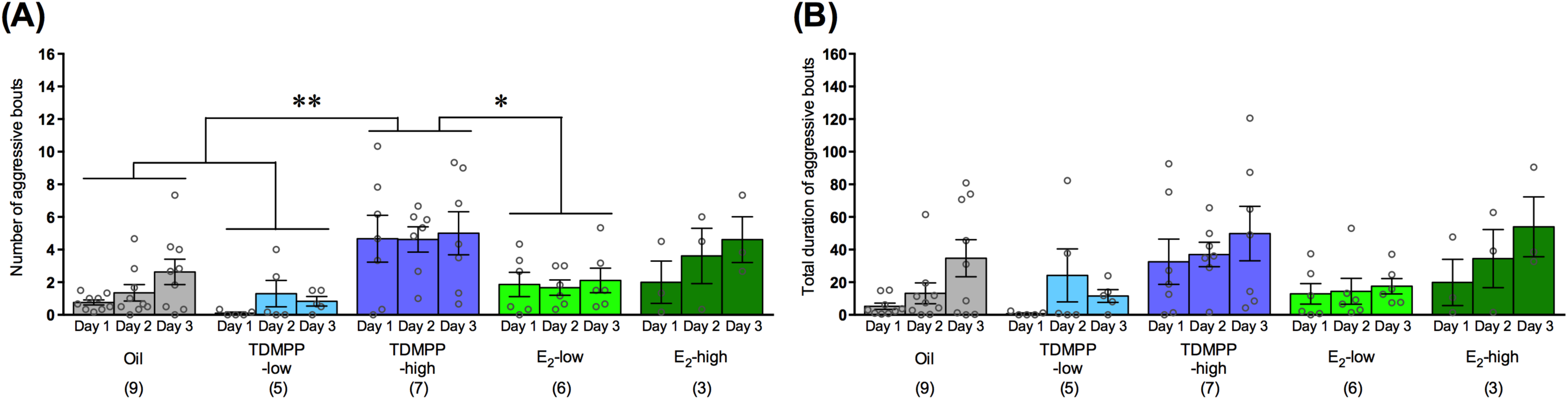
Effect of developmental TDMPP exposure on male aggressive behavior. **(A)** Total number and **(B)** the duration of aggressive bouts toward intruder stimuli. The number in parentheses indicate number of liters. The data are presented as the mean ± SEM. \*\**P* < 0.01, \**P* < 0.05.

### Enzyme immunoassay for plasma testosterone and estradiol

No differences among the male groups were found in plasma testosterone levels [treatment: *F*(4,25) = 1.098, *P* = 0.379, Figure 11A]. No differences in plasma estradiol concentration between the Oil and any other group were found, whereas males in the E_2_-high group showed significantly higher concentrations compared to the TDMPP-low, TDMPP-high, and E_2_-low groups [treatment: *F*(4,25) = 5.725, *P* = 0.002; Bonferroni *post hoc* test: *P* = 0.003, E_2_-high vs. TDMPP-high or E_2_-low; *P* = 0.027, E_2_-high vs. TDMPP-low; Figure11B].

**Figure 11:**
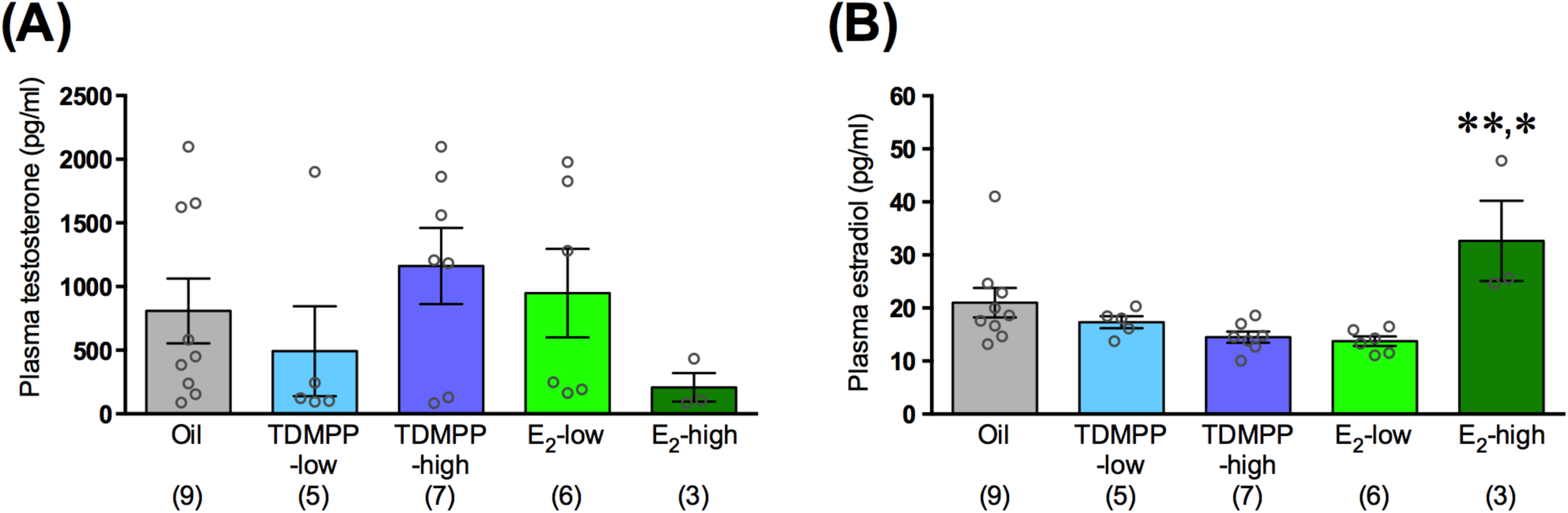
Effect of developmental TDMPP exposure on plasma testosterone and estradiol concentration in male. Plasma concentration of **(A)** testosterone and **(B)** estradiol. The number in parentheses indicate number of liters. The data are presented as the mean ± SEM. \*\**P* < 0.01 vs TDMPP-high, or E_2_-low, \**P* < 0.05 vs TDMPP-low.

### Female sexual behavior

The lordosis quotient, an index of sexual receptivity, in the mice of the TDMPP-high, E_2_-low and E_2_-high groups was significantly reduced compared to that of the Oil group [treatment: *F*(4,24) = 6.822, *P* = 0.001; Bonferroni *post hoc* test, *P* = 0.001, TDMPP-high vs. Oil; *P* = 0.008, E_2_-low vs. Oil; *P* = 0.043, E_2_-high vs. Oil; treatment × day: *F*(16,96) = 1.228, *P* = 0.261; Figure 12], demonstrating that the endocrine-disrupting action of the exposure to either E_2_ or TDMPP during the critical period of brain sexual differentiation impairs the development of sexual receptive behavior.

**Figure 12:**
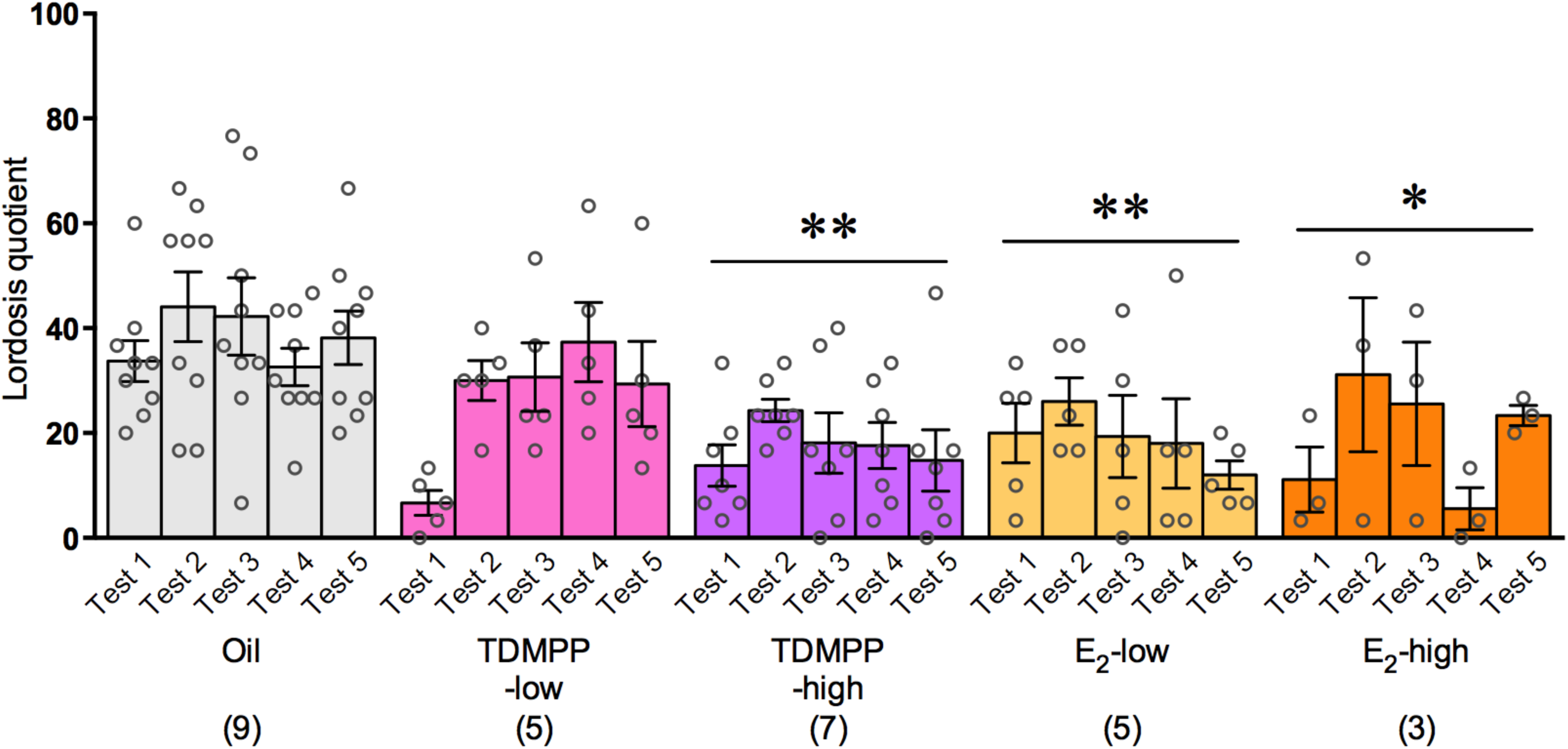
Effect of developmental TDMPP exposure on female sexual behavior. The number in parentheses indicate number of liters. The data are presented as the mean ± SEM. \*\**P* < 0.01, \**P* < 0.05 vs Oil.

### Analysis of sexually dimorphic nuclei

Calb-SDN volume and cell number were significantly higher in the male than in the female Oil group (Student t-test: *P* < 0.001, Figure 13A, B, C, and D). In females, Calb-SDN volume and cell number in the TDMPP-low, TDMPP-high and E_2_-low groups were significantly higher than in the Oil group (ANOVA: volume: *F*(3,15) = 22.734, *P* < 0.001; Bonferroni *post hoc* test: *P* < 0.001, TDMPP-low and TDMPP-high vs. Oil, P = 0.001, E_2_-low vs. Oil, number: *F*(3,15) = 19.012, *P* < 0.001; Bonferroni *post hoc* test: *P* < 0.001, TDMPP-low, TDMPP-high and E_2_-low vs. Oil, Figure 13B and D). In males, Calb-SDN volume and cell number of the TDMPP-low group were significantly higher than the Oil group, but there was no significant difference among the Oil, TDMPP-high and E_2_-low groups (ANOVA: volume: *F*(3,14) = 14.372, *P* < 0.001; Bonferroni *post hoc* test: *P* < 0.001, TDMPP-low vs. Oil, number: *F*(3,14) = 4.678, *P* = 0.018; Bonferroni *post hoc* test: *P* = 0.021, TDMPP-low vs. Oil Figure 13A and C).

**Figure 13:**
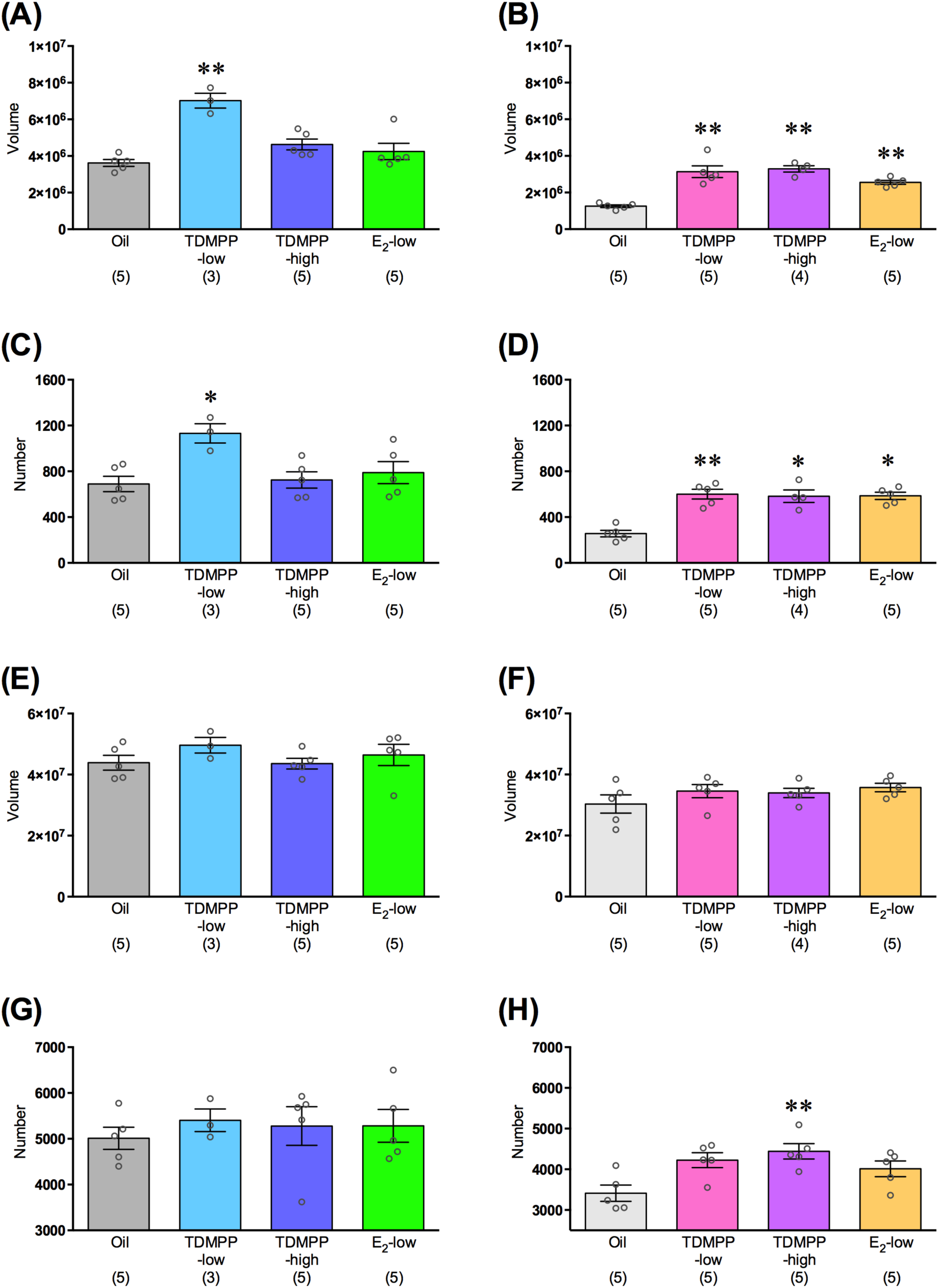
Effect of developmental TDMPP exposure on the sexually dimorphic nuclei. **(A,C,E,G)** male and **(B,D,F,H)** female of each treatment group. (A,B) The volume and (C,D) number of cells in the Calb-SDN. (E,F) The volume and (G,H) number of cells in the BNSTp. The number in parentheses indicate number of animals. The data are presented as the mean ± SEM. \*\**P* < 0.01, \**P* < 0.05 vs Oil.

Similarly, BNSTp volume and cell number were significantly higher in the male than in the female Oil group (Student t-test: *P* = 0.001, Figure 13E,F,G and H). BNSTp cell number in the females of the TDMPP-high groups were significantly higher than in the Oil group (ANOVA: *F*(3,15) = 5.369, *P* = 0.009; Bonferroni *post hoc* test: *P* = 0.009, TDMPP-high vs. Oil, Figure 13H), whereas no difference was found in the volume (ANOVA: *F*(3,16) = 1.221, *P* = 0.334, Figure 13F). In males, there was no difference in either BNSTp volume or cell number between groups (ANOVA: volume: *F*(3,14) = 0.871, *P* = 0.480, number: *F*(3,14) = 0.216, *P* = 0.884, Figure 13E and G).

### Concentrations of TDMPP and its metabolites in brain

Using MS and MS/MS, we first detected TDMPP and its metabolites in neonatal (PND 1) brain samples 16 h after treatment with TDMPP, based on precise molecular weight information about TDMPP and its metabolites and related ions. We detected TDMPP and 4 different metabolites, denoted by di(2,6-dimethylphenyl) phosphate (DDMPP), TDMPP-M1, TDMPP-M2-1, and TDMPP-M2-2. From the molecular structure of such metabolites, a number of conclusions could be drawn: (1) DDMPP is a hydroxylation metabolite of TDMPP; (2) TDMPP-M2-1 and TDMPP-M2-2 are oxidation metabolites of TDMPP; and (3) TDMPP-M1 is an oxidation metabolite of DDMPP, or a hydroxylation metabolite of TDMPP-M2-1 or TDMPP-M2-2 (Supplemental figure 1). We measured the levels of TDMPP and its metabolites in fetal (GD 16) and neonatal (PND 1) brain samples at 0, 8, 16, 24 and 48 h after treatment with TDMPP. In the case of fetal brain samples, TDMPP levels rose to 70 ng/g 8 h after treatment, corresponding to 0.000070% of the treatment dose (5,000 µg). In the case of neonatal brain samples, TDMPP and its metabolites were detected 8 h after treatment, and the TDMPP level rose to 870 ng/g 16 h after treatment, corresponding to 0.017% of the treatment dose (500 µg), suggesting that the relative accumulation of TDMPP is higher in the neonatal than in the fetal brain. In the fetal brain, TDMPP was also still detectable at 16, 24, and 48 h, and its levels were 34%, 93%, and 28% of the 8-hour level, respectively. In the neonatal brain, TDMPP was still detectable at 24 and 48 h, with levels corresponding to 27% and 16% of that at 16 h, respectively (Supplemental table 1 and 2).

## Discussion

In this study, we evaluated the endocrine-disrupting action of TDMPP, an impurity in flame retardant formulations of PBDMPP not recognized as an endocrine disruptor, although the acute and chronic toxicities of TDMPP have already been evaluated (Van der Veen & de Boer, 2012). However, a recent *in vitro* screening of representative phosphate flame retardants and related compounds found the activation of ER-α by TDMPP to be at a level corresponding to about 1/10,000 of that of estradiol. This activation is the highest among 23 chemicals examined in the study, and the activation potency of TDMPP is considered to be similar to that of bisphenol-A, reported in previous literature (Suzuki et al., 2013). Thus, we judged that the endocrine-disrupting action of TDMPP exposure should be preferentially examined compared to other phosphate flame retardants and related chemicals. To evaluate the estrogenic activity of TDMPP, we decided to use a mouse model to specifically investigate the impairment in brain sexual differentiation. The influence of chemicals having estrogenic activity, such as diethylstilbestrol, bisphenol-A, and a certain type of flavonoids, is known to disturb the brain sexual differentiation when administered during either late pregnancy or the neonatal period, i.e. the critical periods of brain differentiation in rodents (Mendes, 2002; Vandenberg et al., 2009). Therefore, we administered TDMPP, and E_2_ as a positive control, to mice throughout the critical period and examined the behavioral, physiological and histological changes related to reproduction.

Given that no significant differences in body weight and AGD during development in the TDMPP-treated groups were found, gross morphological effects could not be detected, at least with the treatment schedule we used. However, we cannot exclude the possibility that TDMPP affects the morphogenesis of estrogen-sensitive organs, because it has been reported that estrogenic agents in general affect AGD when administered at earlier time points (Honma et al., 2002). On the other hand, we could detect impairments in different aspects of female physiology and behavior due to developmental exposure to TDMPP, although the open field and light-dark transition tests were not altered. Acceleration of vaginal opening, an index of precocious puberty, and irregular estrous cycle, an index of impairment in the hypothalamo-pituitary-gonadal axis, are known to be typical effects of exposure to estrogenic chemicals of rodents in the critical period of brain sexual differentiation: Our results obtained in both TDMPP-and E_2_-treated females were in agreement with the reported impairments in animals developmentally exposed to estrogenic chemicals. Similarly, sexual receptivity in females, examined by the lordosis behavior test, was impaired by developmental exposure to TDMPP. Since these physiological and behavioral changes mostly coincided with those due to exposure to the positive control E_2_, we conclude that the effect of TDMPP on female reproductive function impairment is mediated by the activation of the estrogen signaling pathways, as expected.

In rodents, the differentiation of core sexual behavior and reproductive physiology is thought to be influenced by exposure to gonadal hormones, whereas the sexual differentiation of various other aspects of physiology and behavior, such as social communication, is thought to be determined at least in part by the interaction of gonadal hormones and sex chromosomes (Arnold, 2004; McCarthy & Arnold, 2011, Maekawa et al., 2014). In particular, sexual and aggressive behaviors are thought to be masculinized and de-feminized by the exposure to testosterone secreted from testes during the critical period of brain sexual differentiation (Bronson & Desjardins, 1968; Pfaff & Zigmond, 1971). Testosterone that enters the brain can be converted into E_2_ by brain aromatase, and the activation of estrogen receptors by locally synthesized brain E_2_ mediates the process of brain masculinization and de-feminization (Naftolin et al., 1971; McEwen et al., 1977). Conversely, the lack of exposure to testosterone and/or estrogen during the critical period of sexual differentiation is essential for the differentiation of female receptive behavior. Indeed, the exogenous injection of either testosterone or estrogen during the critical period of sexual differentiation was reported to impair the development of sexual behavior in females (McDonald & Doughty, 1972; Kouki et al., 2003; Kanaya & Yamanouchi, 2012). Furthermore, since loss-of-function in estrogen signaling by ER-α, but not ER-β,gene knockout impairs masculinization and de-feminization (Ogawa et al., 1998, 1999), the process of differentiation of female receptive behavior and reproductive physiology is predominantly mediated via the ER-α pathway. In order to know whether ER-α in the brain is “directly” involved in the physiological and reproductive functions, we devised various experimental approaches. As a first approach, through chemical exposure measurements, we confirmed the direct transfer of the injected TDMPP and its metabolites to both the fetal and neonatal brain. As a second approach, through histological techniques, we examined the effect of TDMPP on the formation of the typical male-dominant sexual dimorphic nuclei, finding that the volume and cell number of the Calb-SDN and BNSTp were increased up to the level of male mice by the developmental exposure of females to TDMPP. It has been reported that the treatment with an ER-α, but not ER-β, agonist mimics the effect of E_2_ on the establishment of sexual dimorphism in the sexually dimorphic nucleus of the preoptic area, including the Calb-SDN (Patchev et al., 2004; Tsukahara 2009; Sickel & McCarthy, 2000). It has also been reported that the volume and number of neurons in the BNSTp were feminized in male mice deficient in the ER-α gene (Tsukahara et al., 2011). These reports demonstrate that cellular signaling downstream of ER-α is required for the formation of these nuclei. Taken together, these previous reports and our experiments show that the activation of ER-α by TDMPP in the brain could be a most likely mechanism explaining the behavioral and physiological changes in female mice.

Concerning male behavior, the mice in the TDMPP-high group revealed higher aggressive behavior compared to those in the control Oil group, whereas there was no difference in male sexual behavior among groups. This behavioral change might reflect the impact of brain hyper-masculinization by developmental exposure to TDMPP. In terms of hormonal effects, male aggressive behavior is known to be regulated by brain sexual differentiation in the critical period, and sex steroid hormonal levels in the adult (Bronson & Desjardins, 1968). Since testosterone and estradiol levels in the adults of the TDMPP-high groups were similar to those of the Oil control group, brain sexual differentiation during the developmental period might lead to higher aggressiveness. We also examined the volume and cell number of sexual dimorphic nuclei in the male groups, but no significant change was appreciated in either Calb-SDN or BNSTp in the TDMPP-high group, although the volume and cell number of Calb-SDN were rather significantly increased in the TDMPP-low group. Therefore, we cannot find a clear histological change corresponding to the higher aggression shown by the TDMPP-high group. To summarize the results on male mice, the effect of TDMPP was limited compared to females, and the group revealing behavioral abnormality was not consistent with that revealing histological abnormality. Since the male brain is naturally formed under developmental exposure to estradiol converted from testosterone within the brain, the additional estrogenic effect of TDMPP might be less pronounced in terms of activation of ER-α.

The fertility rate of mating pairs in each group was also examined (Supplemental Table 1). Mating pairs of the TDMPP-low group showed low fertility, while all mating pairs of the TDMPP-high group were infertile. These results clearly demonstrate that TDMPP exposure causes lower birth rate, presumably due to the lowered reproductive behavior and physiology in females. Usage of organophosphate flame retardants has recently increased in many household products, and the transfer of organophosphate flame retardants to house dust in living situations has been reported worldwide (Stapleton et al., 2009). Thus, not only industrial but also environmental exposure to TDMPP will be increased with the increasing use of organophosphorus phosphate flame retardants in the future. Based on our empirical data, the contamination of TDMPP in household products, and the subsequent home exposure to TDMPP during pregnancy and/or the neonatal period could be suspected to endanger women’s health in the next generation. Norms limiting the contamination of TDMPP in products are thus required.

## Acknowledgements

This work was supported by National Institute for Environmental Studies, Japan [NIES Research Project (No.130)] to FM.

## Supplementary methods

### Effect of TDMPP on the induction of lordosis and uterine weight in sexually mature females

Female C57BL/6J mice purchased from CLEA Japan (Tokyo, Japan) were ovariectomized at 11 weeks of age under isoflurane anesthesia. At 12 weeks of age (Test 1), either 17β-estradiol (5 µg/2 ml sesame oil), TDMPP (99.9%, Hayashi Pure Chemical Ltd, Osaka, Japan, 50 mg/2 ml sesame oil), or sesame oil (2 ml) were subcutaneously injected twice, at 48 and 24 h before testing. Additionally, progesterone (250 µg/0.1 ml sesame oil) was subcutaneously injected 4 h before testing Females were tested for sexual behavior toward a sexually experienced ICR/JCL male mouse in the male’s home cage. Each test lasted until females received either 15 mounts or 15 intromissions. The number of lordosis responses to either mount or intromission was scored for each mouse. A lordosis quotient was calculated by dividing the number of lordosis responses by 15 mounts or intromission. The same test was also performed at 13 weeks of age (Test 2). Immediately after the last lordosis test, females were sacrificed by decapitation and uterine weight was measured.

### Reproduction rate in mating

Four to five female-male pairs in the Oil, TDMPP-low, TDMPP-high and E_2_-low groups at 15 weeks of age were randomly selected and bred for three estrous cycles to determine whether females showed vaginal plug. In addition, the number of pups was counted if the female became pregnant and delivered.

### Ovarian morphology

Ovaries at 14 weeks of age were fixed by 4% paraformaldehyde in 0.05 M PBS, embedded in paraffin blocks and cut by microtome at a thickness of 10 ?m. Sections were stained by conventional hematoxylin-eosin staining and were observed by light microscopy.

## Supplementary results

### Body weight

No significant effect of TDMPP exposure was found on the body weight of either male or female mice at birth, PND 21 and 10 weeks of age (Figure 2A-F). On the other hand, a slight but significant difference was found among the E_2_ positive control groups.

In male mice, there was no difference between the Oil and the other groups, whereas there was a statistically significant difference at birth between the E_2_-low and E_2_-high groups [*F*(4,25) = 5.215, *P* = 0.003; Bonferroni *post hoc* test: *P* = 0.002, E_2_-low vs. E_2_-high; Figure 2A]. This difference between male groups disappeared at PND 21 and 10 weeks of age [PND 21: *F*(4,25) = 1.451, *P* = 0.247, Figure 2B; 10 weeks: *F*(4,25) = 1.154, *P* = 0.355, Figure 2C]. Additionally, female mice in the E_2_-high group were significantly heavier than those in the E_2_-low and TDMPP-low groups [*F*(4,24) = 5.526, *P* = 0.003; Bonferroni *post hoc* test: *P* = 0.007, E_2_-high vs. E_2_-low or P = 0.004, E_2_-high vs. TDMPP-low; Figure 2D], whereas there was no difference between the Oil group and other groups. At PND 21, there was no difference in their body weight [*F*(4,24) = 1.097, *P* = 0.381, Figure 2E]. At 10 weeks of age, although ANOVA detected a significant treatment effect, no differences between groups were found in the Bonferroni *post hoc* test [*F*(4,24) = 3.112, *P* = 0.034; Bonferroni *post hoc* test, ns; Figure 2F].

### Anogenital distance

In male mice, although ANOVA detected a significant treatment effect on AGD at birth, no differences between groups were revealed by Bonferroni *post hoc* test [*F*(4,25) = 3.508, *P* = 0.021; Bonferroni *post hoc* test, ns; Figure 3A]. At PND 21, AGD did not differ between groups in males [*F*(4,25) = 2.065, *P* = 0.116; Figure 3B]. In female mice, no differences were found in AGD among groups at birth or PND 21 [at birth: *F*(4,24) = 1.022, *P* = 0.416, Figure 3C; PND 21: *F*(4,24) = 1.309, *P* = 0.295, Figure 3D].

### Open field test

In males, no difference was found in total moving distance [treatment: *F*(4,25) = 0.689, *P* = 0.606; treatment × day: *F*(4,25) = 2.264, *P* = 0.091; Figure 7A] and center time between groups [treatment: *F*(4,25) = 0.812, *P* = 0.529; treatment × day: *F*(4,25) = 1.278, *P* = 0.305; Figure 7B]. Although female mice in the E_2_-low and E_2_-high groups showed decreased total moving distance compared to the TDMPP-low group, there was no difference between TDMPP-exposed groups and the Oil group [treatment: *F*(4,24) = 3.975, *P* = 0.013; Bonferroni *post hoc* test: *P* = 0.036, E_2_-low vs. TDMPP-low; *P* = 0.020, E_2_-high vs. TDMPP-low; treatment × day: *F*(4,24) = 2.616, *P* = 0.060; Figure 7C]. There was no difference in center time among groups in females [treatment: *F*(4,24) = 1.201, *P* = 0.336; treatment × day: *F*(4,24) = 0.225, *P* = 0.921; Figure 7D].

### Effect of TDMPP on lordosis induction and uterine weight in sexually mature females

The lordosis quotient was significantly increased by treatment with 17β-estradiol in both Test 1 and 2 (treatment: *F*(2,15) = 38.645, *P* < 0.001; Bonferroni *post hoc* test: *P* < 0.001, E_2_ vs. Oil or TDMPP; Supplemental figure 2). Although there was no statistical difference, five out of six female (83.3%) in TMDPP treated group showed at least one lordosis response whereas only one out of six female (16.7%) in oil treated group showed lordosis response in test 2 (Supplemental table 3). As for uterine weight, both estradiol benzoate and 2,6-TDMPP treatments significantly increased uterine weight (*F*(2,15) = 30.331, *P* < 0.001; Bonferroni *post hoc* test: *P* < 0.001, E_2_ or TDMPP vs. Oil; Supplemental figure 3), demonstrating that TDMPP shows estrogenic action also in adults, even if the doses necessary for the effect should be higher than in the perinatal period.

### Reproduction rate in mating

All females of 5 female-male pairs in the control group showed vaginal plugs by the first or second estrous cycle, and 7.8 ± 1.1 pups (mean ± SEM) were delivered from five females. On the other hand, only one out of 5 females in the E_2_-low group showed a vaginal plug, and none of them delivered. Similarly, none of the 5 females in the TDMPP-high group showed a vaginal plug, and no delivery occurred. In the TDMPP-low group 3 out of 4 females showed vaginal plugs within 3 estrous cycles, and deliveries occurred in 3 out of 4 females, although all the pups of a pregnant female were dead (Supplemental table 4).

### Ovarian morphology

In the Oil and TDMPP-low groups, ovarian follicles and the corpus luteum were found. On the other hand, the corpus luteum scarcely appeared in ovaries of TDMPP-high, E_2_-low and E_2_-high groups, indicating that the release of mature eggs from ovaries was impaired (Supplemental figure 4).

## Supplemental figure legends

**Supplemental Figure 1: The molecular structures of PBDMPP, TDMPP and its 4 different metabolites.**

**Supplemental Figure 2: Effect of TDMPP administration on lordosis induction.** The number in parentheses indicate number of animals. The data are presented as the mean ± SEM. \*\**P* < 0.01.

**Supplemental Figure 3: Examination of uterotropic property of TDMPP.** The number in parentheses indicate number of animals. The data are presented as the mean ± SEM. \*\**P* < 0.01 vs. Oil.

**Supplemental Figure 4: Effect of TDMPP administration on ovarian morphorogy.** Representative photomicrographs of ovary sections from females of each treatment group. Scale bar indicate 200 µm.

## References

Alonso-Magdalena, P., Ropero, A. B., Soriano, S., García-Arévalo, M., Ripoll, C., Fuentes, E.,… – Nadal, Á. (2012). Bisphenol-A acts as a potent estrogen via non-classical estrogen triggered pathways. Molecular and cellular endocrinology, 355(2), 201–207. PMID:22227557; doi:10.1016/j.mce.2011.12.012

Arnold, A. P. (2004). Sex chromosomes and brain gender. Nature Reviews Neuroscience, 5(9), 701. PMID:15322528; doi:10.1038/nrn1494

Bronson, F. H., – Desjardins, C. (1968). Aggression in adult mice: Modification by neonatal injections of gonadal hormones. Science, 161(3842), 705–706. PMID:5691022

Büdefeld T, Grgurevic N, Tobet SA, Majdic G. Sex differences in brain developing in the presence or absence of gonads. Dev Neurobiol. 2008 Jun;68(7):981–95. PMID:18418875; doi:10.1002/dneu.20638.

Cora, M. C., Kooistra, L., – Travlos, G. (2015). Vaginal cytology of the laboratory rat and mouse: review and criteria for the staging of the estrous cycle using stained vaginal smears. Toxicologic pathology, 43(6), 776–793. PMID:25739587; doi:10.1177/0192623315570339

Diamanti-Kandarakis, E., Bourguignon, J. P., Giudice, L. C., Hauser, R., Prins, G. S., Soto, A. M.,… – Gore, A. C. (2009). Endocrine-disrupting chemicals: an Endocrine Society scientific statement. Endocrine reviews, 30(4), 293–342. PMID:19502515; doi:10.1210/er.2009-0002

European Parliament (2002). Directive 2002/95/EC of the European Parliament and of the Council on the restriction of the use of certain hazardous substances in electrical and electronic equipment Off. J. Eur. Union L 2003, 37, 19–23.

Gilmore, R. F., Varnum, M. M., – Forger, N. G. (2012). Effects of blocking developmental cell death on sexually dimorphic calbindin cell groups in the preoptic area and bed nucleus of the stria terminalis. Biology of sex differences, 3(1), 5. PMID:22336348; doi:10.1186/2042-6410-3-5

Gould, J. C., Leonard, L. S., Maness, S. C., Wagner, B. L., Conner, K., Zacharewski, T.,… – Gaido, K. W. (1998). Bisphenol A interacts with the estrogen receptor α in a distinct manner from estradiol. Molecular and cellular endocrinology, 142(1-2), 203–214. PMID:9783916

Honma, S., Suzuki, A., Buchanan, D. L., Katsu, Y., Watanabe, H., – Iguchi, T. (2002). Low dose effect of in utero exposure to bisphenol A and diethylstilbestrol on female mouse reproduction. Reproductive Toxicology, 16(2), 117–122. PMID:11955942

Kanaya, M., – Yamanouchi, K. (2012). Defeminization of brain functions by a single injection of estrogen receptor α or β agonist in neonatal female rats. Neuroendocrinology, 95(4), 297–304. PMID:22327340; doi:10.1159/000332128

Kouki, T., Kishitake, M., Okamoto, M., Oosuka, I., Takebe, M., – Yamanouchi, K. (2003). Effects of neonatal treatment with phytoestrogens, genistein and daidzein, on sex difference in female rat brain function: estrous cycle and lordosis. Hormones and Behavior, 44(2), 140–145. PMID:13129486

Langer, A., Meleis, A., Knaul, F. M., Atun, R., Aran, M., Arreola-Ornelas, H.,… – Claeson, M. (2015). Women and health: the key for sustainable development. The Lancet, 386(9999), 1165–1210. PMID:26051370; doi:10.1016/S0140-6736(15)60497-4

MacLusky, N. J., and Naftolin, F. (1981). Sexual differentiation of the central nervous system. Science 211, 1294–1302. PMID:6163211; doi:10.1126/science.6163211

Maekawa, F., Tsukahara, S., Kawashima, T., Nohara, K., – Ohki-Hamazaki, H. (2014). The mechanisms underlying sexual differentiation of behavior and physiology in mammals and birds: relative contributions of sex steroids and sex chromosomes. Frontiers in neuroscience, 8, 242. PMID:25177264; doi:10.3389/fnins.2014.00242

Matsukami, H., Tue, N.M., Suzuki, G., Someya, M., Tuyen, le H., Viet, P.H., Takahashi, S., Tanabe, S., Takigami, H. Flame retardant emission from e-waste recycling operation in northern Vietnam: environmental occurrence of emerging organophosphorus esters used as alternatives for PBDEs. Sci Total Environ. 2015 May 1;514:492–9. doi:10.1016/j.scitotenv.2015.02.008

McCarthy, M. M., – Arnold, A. P. (2011). Reframing sexual differentiation of the brain. Nature neuroscience, 14(6), 677. PMID:21613996; doi:10.1038/nn.2834

McDonald, P. G., – Doughty, C. (1972). Comparison of the effect of neonatal administration of testosterone and dihydrotestosterone in the female rat. Journal of reproduction and fertility, 30(1), 55–62. PMID:5064322

McEwen, B. S., Lieberburg, I., Chaptal, C., – Krey, L. C. (1977). Aromatization: important for sexual differentiation of the neonatal rat brain. Hormones and behavior, 9(3), 249–263. PMID:611076

Meerts, I. A., Letcher, R. J., Hoving, S., Marsh, G., Bergman, A., Lemmen, J. G.,… – Brouwer, A. (2001). In vitro estrogenicity of polybrominated diphenyl ethers, hydroxylated PDBEs, and polybrominated bisphenol A compounds. Environmental health perspectives, 109(4), 399. PMID:11335189; doi:10.1289/ehp.01109399

Mendes, J. A. (2002). The endocrine disrupters: a major medical challenge. Food and Chemical Toxicology, 40(6), 781–788. PMID:11983272

Moe, Y., Tanaka, T., Morishita, M., Ohata, R., Nakahara, C., Kawashima, T.,… – Tsukahara, S. (2016). A comparative study of sex difference in calbindin neurons among mice, musk shrews, and Japanese quails. Neuroscience letters, 631, 63–69. PMID:27531632; doi:10.1016/j.neulet.2016.08.018

Naftolin, F., Ryan, K. J., – Petro, Z. (1971). Aromatization of androstenedione by the diencephalon. The Journal of Clinical Endocrinology – Metabolism, 33(2), 368–370. PMID:4935642; doi:10.1210/jcem-33-2-368

Ogawa, S., Chan, J., Chester, A. E., Gustafsson, J. Å., Korach, K. S., – Pfaff, D. W. (1999). Survival of reproductive behaviors in estrogen receptor β gene-deficient (βERKO) male and female mice. Proceedings of the National Academy of Sciences, 96(22), 12887–12892. PMID:10536018

Ogawa, S., Washburn, T. F., Taylor, J., Lubahn, D. B., Korach, K. S., – Pfaff, D. W. (1998). Modifications of testosterone-dependent behaviors by estrogen receptor-α gene disruption in male mice. Endocrinology, 139(12), 5058–5069. PMID:9832445; doi:10.1210/endo.139.12.6358

Orikasa C, Sakuma Y (2010) Estrogen configures the sexual dimorphism in the preoptic area of C57/BL6J and ddN strains of mice. Journal of Comparative Neurology 518(17): 3618–3629. PMID:20593361; doi:10.1002/cne.22419

Patchev, A. V., Gotz, F., and Rohde, W. (2004). Differential role of estrogen receptor isoforms in sex-specific brain organization. FASEB J. 18, 1568–1570. PMID:15289439; doi:10.1096/fj.04-1959fje

Paxinos, G., – Franklin, K. B. (2004). The mouse brain in stereotaxic coordinates. Gulf professional publishing.

Pfaff, D. W., – Zigmond, R. E. (1971). Neonatal androgen effects on sexual and non-sexual behavior of adult rats tested under various hormone regimes. Neuroendocrinology, 7(3), 129–145. PMID:5546026; doi:10.1159/000121961

Phoenix, C. H., Goy, R. W., Gerall, A. A., and Young, W. C. (1959). Organizing action of prenatally administered testosterone propionate on the tissues mediating mating behavior in the female guinea pig. Endocrinology 65, 369–382. PMID:14432658; doi:10.1210/endo-65-3-369

Prendergast, B. J., Onishi, K. G., – Zucker, I. (2014). Female mice liberated for inclusion in neuroscience and biomedical research. Neuroscience – Biobehavioral Reviews, 40, 1–5. PMID:24456941; doi:10.1016/j.neubiorev.2014.01.001

Sano, K., Isobe, T., Yang, J., Win-Shwe, T. T., Yoshikane, M., Nakayama, S. F.,… – Tohyama, C. (2016). In utero and lactational exposure to acetamiprid induces abnormalities in socio-sexual and anxiety-related behaviors of male mice. Frontiers in neuroscience, 10, 228. MID:27375407; doi:10.3389/fnins.2016.00228

Sickel, M. J., – McCarthy, M. M. (2000). Calbindin-D28k immunoreactivity is a marker for a subdivision of the sexually dimorphic nucleus of the preoptic area of the rat: developmental profile and gonadal steroid modulation. Journal of neuroendocrinology, 12(5), 397–402. PMID:10792577

Stapleton, H. M., Klosterhaus, S., Eagle, S., Fuh, J., Meeker, J. D., Blum, A., – Webster, T. F. (2009). Detection of organophosphate flame retardants in furniture foam and US house dust. Environmental science – technology, 43(19), 7490–7495. PMID:19848166

Suzuki, G., Tue, N. M., Malarvannan, G., Sudaryanto, A., Takahashi, S., Tanabe, S.,… – Takigami, H. (2013). Similarities in the endocrine-disrupting potencies of indoor dust and flame retardants by using human osteosarcoma (U2OS) cell-based reporter gene assays. Environmental science – technology, 47(6), 2898–2908. PMID:23398518; doi:10.1021/es304691a

Tsukahara, S. (2009). Sex differences and the roles of sex steroids in apoptosis of sexually dimorphic nuclei of the preoptic area in postnatal rats. Journal of neuroendocrinology, 21(4), 370–376. PMID:19226350; doi:10.1111/j.1365-2826.2009.01855.x

Tsukahara, S., Tsuda, M. C., Kurihara, R., Kato, Y., Kuroda, Y., Nakata, M., et al. (2011). Effects of aromatase or estrogen receptor gene deletion on masculinization of the principal nucleus of the bed nucleus of the stria terminalis of mice. Neuroendocrinology 94, 137–147. PMID:21525731; doi:10.1159/000327541

United Nations Statistical Commission (2016). Report of the Inter-Agency and Expert Group on Sustainable Development Goal Indicators. 2016. [cited Nov 22 2016]. Available from: https://www.unstatsunorg/unsd/statcom/47th-session//2016-2-IAEG-SDGs-Rev1-Epdf

Vandenberg, L. N., Maffini, M. V., Sonnenschein, C., Rubin, B. S., – Soto, A. M. (2009). Bisphenol-A and the great divide: a review of controversies in the field of endocrine disruption. Endocrine reviews, 30(1), 75–95. PMID:19074586; doi:10.1210/er.2008-0021

Van der Veen, I., – de Boer, J. (2012). Phosphorus flame retardants: properties, production, environmental occurrence, toxicity and analysis. Chemosphere, 88(10), 1119–1153. PMID:22537891; doi:10.1016/j.chemosphere.2012.03.067

Wittmann, W., – McLennan, I. S. (2013). The bed nucleus of the stria terminalis has developmental and adult forms in mice, with the male bias in the developmental form being dependent on testicular AMH. Hormones and behavior, 64(4), 605–610. PMID:24012942; doi:10.1016/j.yhbeh.2013.08.017

Zhou, T., Taylor, M. M., DeVito, M. J., – Crofton, K. M. (2002). Developmental exposure to brominated diphenyl ethers results in thyroid hormone disruption. Toxicological Sciences, 66(1), 105–116. PMID:11861977

