## Supplemental Table1-2 for "Estrogenic action by tris(2,6-dimethylphenyl) phosphate, an impurity in resorcinol bis[di(2,6-dimethylphenyl) phosphate] flame retardant formulations, impairs the development of female reproductive functions"

Supplemental table 1. Concentrations (ng/g) of TDMPP and its metabolites in neonatal brain samples.

|  | TDMPP | DDMPP | TDMPP-M1 | TDMPP-M2-1 | TDMPP-M2-2 |
| --- | --- | --- | --- | --- | --- |
| 0hr-M1 | <3 | <3 | <3 | <3 | <3 |
| 0hr-M2 | 170 | <3 | <3 | <3 | <3 |
| 0hr-M3 | <3 | <3 | <3 | <3 | <3 |
| 0hr-F1 | <3 | <3 | <3 | <3 | <3 |
| 0hr-F2 | 4.5 | <3 | <3 | <3 | <3 |
| 0hr-F3 | 83 | <3 | <3 | <3 | <3 |
| 4hr-M1 | 89 | 16 | 10 | <3 | <3 |
| 4hr-M2 | 120 | 13 | <3 | <3 | <3 |
| 4hr-M3 | 120 | 16 | <3 | <3 | <3 |
| 4hr-F1 | 500 | 18 | <3 | <3 | <3 |
| 4hr-F2 | 95 | 20 | <3 | <3 | <3 |
| 4hr-F3 | 120 | <3 | <3 | <3 | <3 |
| 8hr-M1 | 210 | 44 | 24 | <3 | 5.3 |
| 8hr-M2 | 240 | 44 | 24 | <3 | 4.3 |
| 8hr-M3 | 150 | 26 | 18 | <3 | 4.0 |
| 8hr-F1 | 380 | 70 | 30 | 4.5 | 8.0 |
| 8hr-F2 | 250 | 49 | 29 | 3.7 | 8.2 |
| 8hr-F3 | 140 | 29 | 16 | <3 | 4.1 |
| 16hr-M1 | 460 | 110 | 68 | 5.2 | 14 |
| 16hr-M2 | 240 | 60 | 48 | <3 | 6.3 |
| 16hr-M3 | 2900 | 180 | 100 | 9.9 | 27 |
| 16hr-F1 | 190 | 64 | 42 | <3 | 6.4 |
| 16hr-F2 | 220 | 78 | 50 | <3 | 7.6 |
| 16hr-F3 | 1200 | 58 | 38 | <3 | 8.2 |
| 24hr-M1 | 190 | 45 | 27 | <3 | 4.8 |
| 24hr-M2 | 220 | 39 | 26 | <3 | 5.3 |
| 24hr-M3 | 130 | 45 | 32 | <3 | 3.6 |
| 24hr-F1 | 470 | 69 | 53 | 4.5 | 12 |
| 24hr-F2 | 280 | 79 | 48 | <3 | 6.0 |
| 24hr-F3 | 110 | 21 | 13 | <3 | 3.0 |
| 48hr-M1 | 170 | 24 | 13 | <3 | <3 |
| 48hr-M2 | 100 | 20 | 11 | <3 | <3 |
| 48hr-M3 | 160 | 15 | 10 | <3 | <3 |
| 48hr-F1 | 120 | 17 | 13 | <3 | <3 |
| 48hr-F2 | 110 | 21 | 11 | <3 | <3 |
| 48hr-F3 | 160 | 21 | 11 | <3 | <3 |

Supplemental table 2. Concentrations (ng/g) of TDMPP and its metabolites in fetal brain samples.

|  | TDMPP | DDMPP | TDMPP-M1 | TDMPP-M2-1 | TDMPP-M2-2 |
| --- | --- | --- | --- | --- | --- |
| 0hr-1-M1 | <5 | <5 | <5 | <5 | <5 |
| 0hr-1-M2 | <5 | <5 | <5 | <5 | <5 |
| 0hr-1-F1 | <5 | <5 | <5 | <5 | <5 |
| 0hr-1-F2 | <5 | <5 | <5 | <5 | <5 |
| 0hr-2-M1 | <5 | <5 | <5 | <5 | <5 |
| 0hr-2-M2 | <5 | <5 | <5 | <5 | <5 |
| 0hr-2-F1 | <5 | <5 | <5 | <5 | <5 |
| 0hr-2-F2 | <5 | <5 | <5 | <5 | <5 |
| 0hr-3-M1 | 9.2 | <5 | <5 | <5 | <5 |
| 0hr-3-M2 | <5 | <5 | <5 | <5 | <5 |
| 0hr-3-F1 | 11 | <5 | <5 | <5 | <5 |
| 0hr-3-F2 | <5 | <5 | <5 | <5 | <5 |
| 8hr-1-M1 | 7.4 | <5 | <5 | <5 | <5 |
| 8hr-1-M2 | 8.6 | <5 | <5 | <5 | <5 |
| 8hr-1-F1 | 6.8 | <5 | <5 | <5 | <5 |
| 8hr-1-F2 | 8.0 | <5 | <5 | <5 | <5 |
| 8hr-2-M1 | 200 | 69 | 43 | <5 | <5 |
| 8hr-2-M2 | 110 | 75 | 45 | <5 | <5 |
| 8hr-2-F1 | 120 | 60 | 41 | <5 | <5 |
| 8hr-2-F2 | 110 | 65 | 36 | <5 | <5 |
| 8hr-3-M1 | 63 | <5 | <5 | <5 | <5 |
| 8hr-3-M2 | 33 | <5 | <5 | <5 | <5 |
| 8hr-3-F1 | 69 | <5 | <5 | <5 | <5 |
| 8hr-3-F2 | 100 | <5 | <5 | <5 | <5 |
| 16hr-1-M1 | 38 | 52 | 32 | <5 | <5 |
| 16hr-1-M2 | 42 | 64 | 39 | <5 | <5 |
| 16hr-1-F1 | 62 | 58 | 32 | <5 | <5 |
| 16hr-1-M3 | 33 | 47 | 32 | <5 | <5 |
| 16hr-2-M1 | 6.8 | <5 | <5 | <5 | <5 |
| 16hr-2-M2 | 7.2 | <5 | <5 | <5 | <5 |
| 16hr-2-F1 | 7.0 | <5 | <5 | <5 | <5 |
| 16hr-2-F2 | 6.0 | <5 | <5 | <5 | <5 |
| 16hr-3-M1 | 37 | <5 | <5 | <5 | <5 |
| 16hr-3-M2 | 13 | <5 | <5 | <5 | <5 |
| 16hr-3-F1 | 19 | <5 | <5 | <5 | <5 |
| 16hr-3-F2 | 10 | <5 | <5 | <5 | <5 |

Supplemental table 2. (Continued)

|  | TDMPP | DDMPP | TDMPP-M1 | TDMPP-M2-1 | TDMPP-M2-2 |
| --- | --- | --- | --- | --- | --- |
| 24hr-1-M1 | 23 | <5 | <5 | <5 | <5 |
| 24hr-1-M2 | 15 | <5 | <5 | <5 | <5 |
| 24hr-1-F1 | 140 | <5 | <5 | <5 | <5 |
| 24hr-1-F2 | 22 | <5 | <5 | <5 | <5 |
| 24hr-2-M1 | 84 | <5 | <5 | <5 | <5 |
| 24hr-2-M2 | 93 | <5 | 15 | <5 | <5 |
| 24hr-2-F1 | 200 | 15 | <5 | <5 | <5 |
| 24hr-2-F2 | 88 | 21 | <5 | <5 | <5 |
| 24hr-3-M1 | 17 | <5 | <5 | <5 | <5 |
| 24hr-3-M2 | 18 | <5 | <5 | <5 | <5 |
| 24hr-3-F1 | 17 | <5 | <5 | <5 | <5 |
| 24hr-3-F2 | 18 | <5 | <5 | <5 | <5 |
| 48hr-1-M1 | 7.0 | 21 | 13 | <5 | <5 |
| 48hr-1-M2 | 10 | 28 | <5 | <5 | <5 |
| 48hr-1-F1 | 11 | 17 | 14 | <5 | <5 |
| 48hr-1-F2 | 10 | 24 | <5 | <5 | <5 |
| 48hr-2-M1 | 25 | 59 | 31 | <5 | <5 |
| 48hr-2-M2 | 32 | 51 | 23 | <5 | <5 |
| 48hr-2-F1 | 46 | 48 | 28 | <5 | <5 |
| 48hr-2-F2 | 16 | 36 | 28 | <5 | <5 |
| 48hr-3-M1 | 28 | 23 | 14 | <5 | <5 |
| 48hr-3-M2 | 9.7 | 22 | 13 | <5 | <5 |
| 48hr-3-F1 | 20 | 20 | <5 | <5 | <5 |
| 48hr-3-F2 | 16 | 16 | 13 | <5 | <5 |
