## Supplemental Table3 for "Estrogenic action by tris(2,6-dimethylphenyl) phosphate, an impurity in resorcinol bis[di(2,6-dimethylphenyl) phosphate] flame retardant formulations, impairs the development of female reproductive functions"

**Supplemental table 3⏐Number of mice showed lordosis response in each test**

Test 1	Test 2

Oil	1/6 (16.7%)	1/6 (16.7%)

TMDPP	5/6 (83.3%)	6/6 (100%)

E_2_	1/6 (16.7%)	5/6 (83.3%)

Number of mice showed at least one lordosis response in each test; the percentage is given in *parentheses*.
