## Supplemental Table4 for "Estrogenic action by tris(2,6-dimethylphenyl) phosphate, an impurity in resorcinol bis[di(2,6-dimethylphenyl) phosphate] flame retardant formulations, impairs the development of female reproductive functions"

**Supplemental table 4⏐Reproduction rate in mating**

Plug 1	Plug 2	Plug 3	Number of pups

Oil 1	○	4

Oil 2	○	9

Oil 3	○	8

Oil 4	×	○	10

Oil 5	○	6

TMDPP-low 1	○	1

TMDPP-low 2	×	×	×	infertile

TMDPP-low 3	×	×	○	8

TMDPP-low 4	○	all dead

TMDPP-high 1	×	×	×	infertile

TMDPP-high 2	×	×	×	infertile

TMDPP-high 3	×	×	×	infertile

TMDPP-high 4	×	×	×	infertile

TMDPP-high 5	×	×	×	infertile

E_2_-1	×	×	×	infertile

E_2_-2	×	×	×	infertile

E_2_-3	×	×	○	(dam died at GD20)

E_2_-4	×	×	×	infertile

E_2_-5	×	×	×	infertile

○: plug observed and became pregnant.

×: plug observed but did not become pregnant.
