## Supplemental Figure1-4 for "Estrogenic action by tris(2,6-dimethylphenyl) phosphate, an impurity in resorcinol bis[di(2,6-dimethylphenyl) phosphate] flame retardant formulations, impairs the development of female reproductive functions"

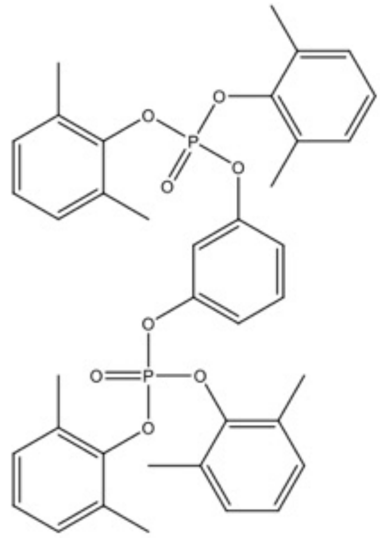

Resorcinol bis[di(2,6-dimethylphenyl) phosphate] (PBDMPP)

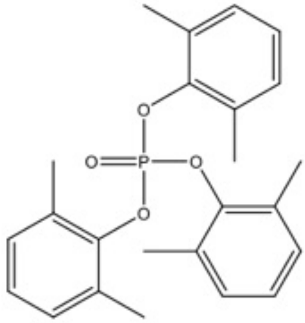

Tris(2,6-dimethylphenyl) phosphate (TDMPP)

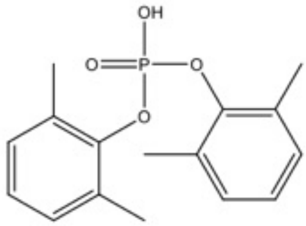

Bis(2,6-dimethylphenyl) phosphate (DDMPP)

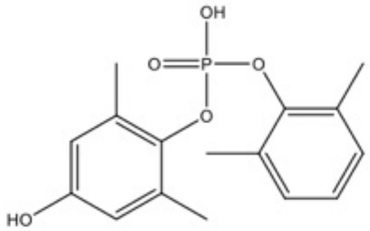

2,6-dimethylphenyl (4-hydroxy-2,6-dimethylphenyl) phosphate (TDMPP-M1)

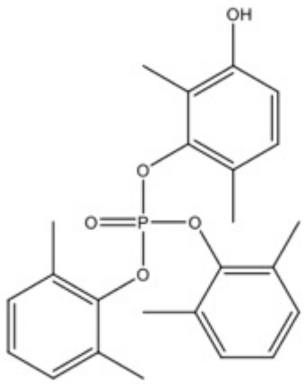

Bis(2,6-dimethylphenyl) (3-hydroxy-2,6-dimethylphenyl) phosphate (TDMPP-M2-1)

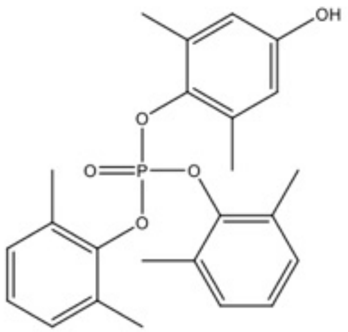

Bis(2,6-dimethylphenyl) (4-hydroxy-2,6-dimethylphenyl) phosphate (TDMPP-M2-2)

**Supplemental figure 1.**

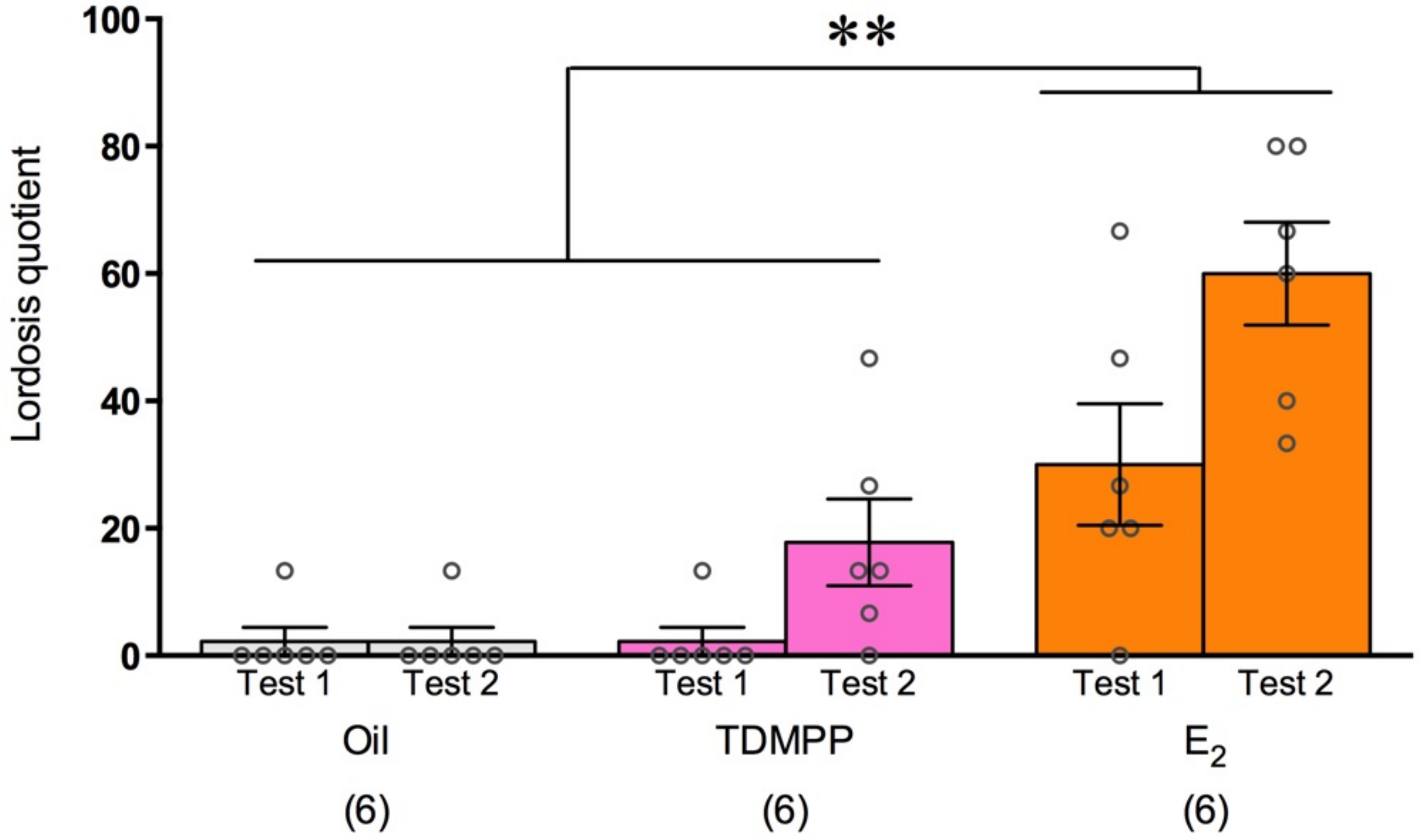


**Supplemental figure 2.**

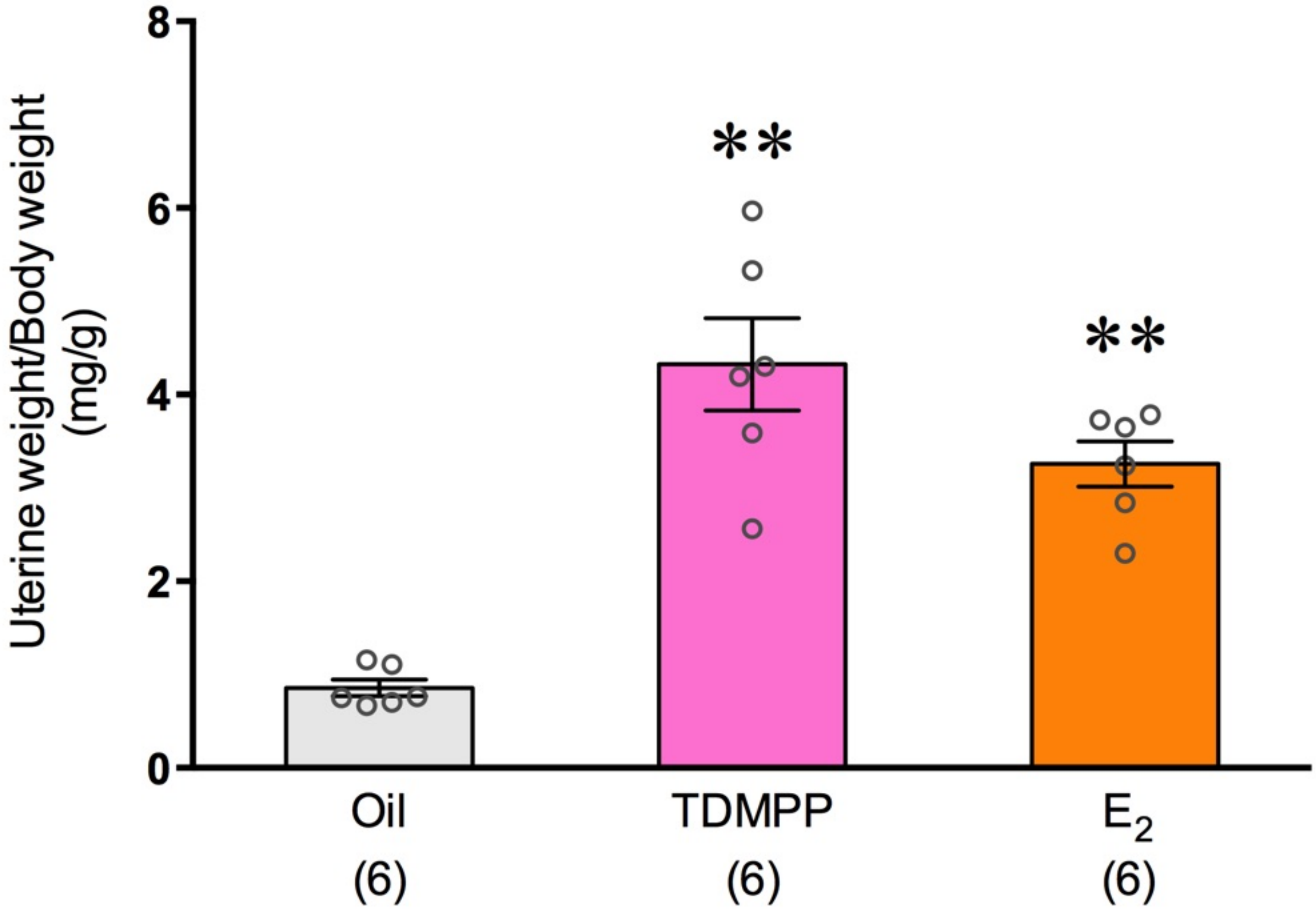


**Supplemental figure 3.**

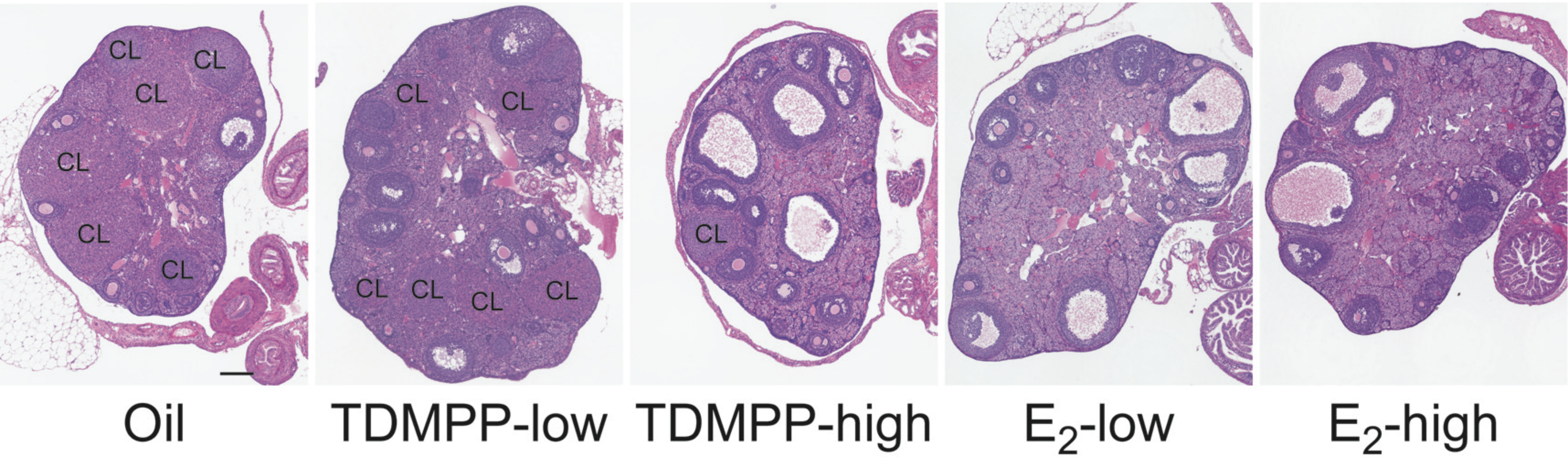


**Supplemental figure 4.**
